# Pharmacologic IRE1 Activation Rewires Cholesterol Metabolism to Suppress T_H_17 Differentiation

**DOI:** 10.64898/2026.11.12.765150

**Authors:** Keishla Sánchez Ortiz, Priyadarshini Chatterjee, William Dion, Meera Pannikkat, Sergei Kutseikin, Taylor Basso, Jie Sun, Prakhyat Mathur, Stanislav Dikiy, Enrique Saez, Jessica E. Thaxton, R. Luke Wiseman, Alejandra Mendoza

## Abstract

T helper 17 (T_H_17) cells are a subset of CD4⁺ T cells that coordinate immune responses against extracellular bacterial and fungal pathogens. Dysregulation of T_H_17 cell responses drives excessive inflammation and contributes to the development of chronic inflammatory and autoimmune diseases, including psoriasis, inflammatory bowel disease, and multiple sclerosis. Here, we demonstrate that pharmacologic activation of the IRE1 signaling arm of the unfolded protein response (UPR) selectively reduces differentiation of T_H_17 cells without impacting the differentiation of other helper T cell subsets. We show that IRE1 activation suppresses T_H_17 differentiation by limiting intracellular cholesterol in differentiating T_H_17 cells through the increased degradation of the rate-limiting cholesterol biosynthesis enzyme HMGCR. Our findings identify IRE1 as a metabolic checkpoint regulating cholesterol biosynthesis in differentiating T_H_17 cells and demonstrate that its pharmacological activation suppresses T_H_17 differentiation, highlighting a potential therapeutic strategy for limiting T_H_17-driven inflammatory diseases.

## INTRODUCTION

CD4⁺ T cells are central orchestrators of adaptive immunity, shaping the quality and magnitude of the immune response through differentiation into specialized effector and memory states^1,2^. Upon priming, naïve CD4⁺ T cells integrate signals from the T cell receptor (TCR), co-stimulatory receptors, and cytokines produced by dendritic cells and other innate immune cells^3^. These cues drive differentiation into distinct lineages, including T_H_1, T_H_2, and T_H_17 cells, as well as peripherally induced regulatory T (T_reg_) cells, which are defined by the expression of lineage-specifying transcription factors T-bet, GATA-3, RORγt, and Foxp3, respectively^2,4–8^. The T_H_1, T_H_2, and T_H_17 CD4^+^ T cell subsets are functionally specialized to orchestrate appropriate responses to distinct pathogen classes, while T_reg_ cells restrain immune activation to preserve tolerance and limit immunopathology^9^. Together, this functional diversity allows CD4⁺ T cells to coordinate and amplify the broader immune response through mechanisms that include the licensing of dendritic cells^10,11^, promotion of phagocytosis^2,12^ and support of CD8⁺ T cell responses^13,14^. Dysregulated or inappropriate activity of each subset is linked to distinct immune-mediated diseases^15–18^, highlighting the need for therapeutic strategies to regulate T effector cell specification and activity.

T_H_17 cells provide essential immunity against bacterial and fungal pathogens^19^ and accumulate at barrier surfaces in response to commensal microbiota such as segmented filamentous bacteria (SFB)^20,21^. However, aberrant T_H_17 responses drive immunopathology in autoimmune and inflammatory diseases, including psoriasis, inflammatory bowel disease, and multiple sclerosis^22–24^. Current strategies to limit T_H_17-driven inflammation rely primarily on antibody-mediated neutralization of cytokines produced by or needed by T_H_17 cells, such as IL-17A^25^ or IL-23^26,27^. However, the effectiveness of these treatments is limited, as a substantial fraction of patients fail to respond or lose response to treatment over time, efficacy varies across T_H_17-associated indications (e.g., IL-17A blockade worsens Crohn’s disease pathology^28,29^) and these agents can predispose patients to mucocutaneous infection^26,28^. These limitations underscore the need for new approaches that reduce T_H_17 differentiation and activity in disease.

Rather than targeting a single cytokine, an alternative strategy to limit T_H_17 cell activity is to exploit the cellular demands of T_H_17 cell differentiation. Differentiation from a naïve CD4⁺ T cell into a cytokine-secreting effector cell imposes substantial metabolic and proteostatic stress due to rapid expansion^30^, the rewiring of glycolytic^31,32^, lipid, and cholesterol biosynthetic pathways^33,34^, and the need to fold and secrete large amounts of cytokines^35^. To adapt effector function to these stresses, immune cells engage stress-responsive signaling pathways such as the endoplasmic reticulum (ER) stress-responsive unfolded protein response (UPR) to regulate immune and tissue homeostasis during the differentiation process^36^.

The UPR comprises three signaling pathways regulated downstream of the ER membrane proteins PERK, ATF6, and IRE1, which together promote transcriptional and translational signaling that adapts cellular physiology following ER insults^37,38^. IRE1, the most evolutionarily conserved arm of the UPR, is activated by ER stress through a mechanism involving autophosphorylation and oligomerization^39,40^, leading to allosteric activation of its cytosolic RNase domain involved in the non-canonical splicing of *Xbp1* mRNA^41–43^. The resulting transcription factor, spliced XBP1 (XBP1s), promotes protective remodeling of cellular physiology by upregulating genes involved in pathways such as ER proteostasis, lipid synthesis, and metabolism^44,45^. Intriguingly, IRE1/XBP1s signaling has been linked to T cell differentiation and function. In CD8⁺ T cells, XBP1s signaling regulates the protein and lipid biosynthetic capacity required for effector expansion and dictates terminal effector versus memory fate^45,46^. Further, chronic IRE1/XBP1s activity within the tumor microenvironment imposes lipid accumulation, metabolic restriction, and loss of cytokine production in tumor-infiltrating lymphocytes^47,48^. In CD4⁺ T cells, IRE1/XBP1s signaling is activated during differentiation^49^, yet IRE1 and XBP1 are dispensable for the specification of the major T effector subsets^49^. Instead, IRE1 activity shapes subset function. For example, pharmacologic or genetic IRE1 inhibition in T_H_2 cells impairs secretion of the lineage-defining cytokines IL-4 and IL-5 and restricts proliferation of activated T_H_2 cells through XBP1s-dependent regulation of cell cycle genes^50,51^. IL-23-driven IRE1 activation has also been linked to enhanced T_H_17 responses in a fungal airway inflammation asthma model^52^. These findings suggest that IRE1 signaling may act as a checkpoint to determine effector T cell fate and that pharmacologically targeting IRE1 activity may offer an opportunity to redirect CD4⁺ T cell specification.

We previously identified the small molecule IXA4 as a selective activator of adaptive IRE1/XBP1s signaling that induces adaptive IRE1/XBP1s signaling without engaging pathologic IRE1 activities such as regulated IRE1-dependent mRNA decay (RIDD) and TRAF2-mediated NF-κB or JNK signaling^53,54^. IXA4 activates IRE1/XBP1s signaling by binding and inhibiting the ER membrane exporter of oxidized glutathione (GSSG), SLC33A1^55^. This leads to the accumulation of GSSG within the ER lumen, which selectively activates IRE1/XBP1s signaling. IXA4 has been widely used to study the biological function and therapeutic implications of IRE1/XBP1s signaling in cellular and *in vivo* models of many diseases, including Charcot-Marie-Tooth Type 1B, obesity-diabetes, amyotrophic lateral sclerosis, and the autoimmune diseases systemic sclerosis and systemic lupus erythematosus^54,56–58^. Thus, IXA4-dependent increases in IRE1 signaling offer a tractable approach to probe how pharmacologic IRE1 activation influences CD4⁺ T cell differentiation.

Here, we show that IXA4-mediated IRE1 activation selectively inhibits *ex vivo* and *in vivo* differentiation of activated CD4⁺ T cells into T_H_17 cells, without affecting differentiation of T_H_1, T_H_2, or T_reg_ cells. We find that IXA4-induced IRE1 activation reduces T_H_17 differentiation by depleting intracellular cholesterol in differentiating T_H_17 cells through a process involving the proteasome-mediated degradation of the rate-limiting cholesterol biosynthesis enzyme 3-hydroxy-3-methylglutaryl-CoA reductase (HMGCR). Supplementation with cholesterol precursors produced downstream of HMGCR restored intracellular cholesterol levels and rescued T_H_17 differentiation in IXA4-treated cells, confirming that IXA4-induced reductions in T_H_17 differentiation are attributed to rewiring of cholesterol metabolism. Finally, we show that treatment of CD4^+^ T cells with the structurally distinct IRE1/XBP1s activator compound IXA62 similarly promotes IRE1-dependent rewiring of cholesterol metabolism to suppress T_H_17 differentiation, further confirming the impact of pharmacologic IRE1 activation on differentiation of this T cell subset. Our results establish a role for IRE1 signaling in regulating cholesterol metabolism during T_H_17 cell differentiation and identify pharmacologic IRE1 activation as a mechanism to selectively limit the differentiation of this pro-inflammatory CD4^+^ T cell lineage.

## RESULTS

### IXA4 decreases differentiation of T_H_17 cells

To assess the impact of our pharmacologic IRE1/XBP1s activator, IXA4, on the *ex vivo* differentiation of CD4⁺ T cell subsets, we isolated naïve CD4⁺ T cells from the spleens of C57BL/6J mice and subjected them to *ex vivo* polarization toward T_H_1, T_H_2, T_H_17, and T_reg_ cell lineages using defined cytokine and antibody conditions (**Fig. 1A; Fig. S1A**)^59^. IXA4 was added 24 h after the polarization cocktail, and cells were allowed to differentiate for an additional 48 h. At 72 h post-polarization, we harvested cells for phenotypic and functional analysis by flow cytometry. We confirmed that IXA4 treatment for 6 h increased expression of the IRE1 target gene *Xbp1s* across all T cell specification conditions (**Fig. 1B**, **Fig. S1B**). Treatment with IXA4 did not influence viability, proliferation, or general activation status, measured by cell surface expression of CD44 and cell numbers, across any of the T cell subsets (**Fig. S1C-F**). Further, IXA4 treatment did not influence expression of lineage-specific transcription factors in the T_H_1 (T-bet), T_H_2 (GATA-3), or T_reg_ (Foxp3) subsets (**Fig. S1G-H**). In contrast, we observed reductions in the expression of the lineage-specifying transcription factor RORγt in CD4^+^ T cells polarized towards the T_H_17 lineage treated with IXA4 (**Fig. 1C-D**). IXA4 treatment during differentiation also reduced production of IL-17A by T_H_17 cells upon restimulation with phorbol 12-myristate 13-acetate (PMA) and ionomycin (**Fig. 1E, Fig. S1I**). We did not observe increased Foxp3 expression in T_H_17 cells differentiating in the presence of IXA4, indicating that IXA4 treatment does not increase T_reg_ differentiation under these conditions (**Fig. S1J**). IXA4 treatment reduced expression of RORγt in differentiating T_H_17 cells in a dose-dependent manner with an EC_50_ of 0.5 µM (**Fig. 1F**), a potency comparable to that observed for IXA4-induced IRE1/XBP1s activation in other cell models^53^. IXA4 treatment also reduced IL-17A production in activated CD4^+^ T cells differentiated in alternative T_H_17 polarizing conditions (IL-23 and IL-1β), indicating that the impact of IXA4 on T_H_17 cell differentiation is independent of the specific polarizing cytokine context (**Fig. S1K**). However, IXA4 treatment does not disrupt expression of RORγt or IL-17A production in cells that had been polarized for 72 h in T_H_17 specification conditions prior to IXA4 treatment (**Fig. S1L**). Collectively, these results indicate that treatment with the IRE1/XBP1s activator IXA4 selectively impairs the *ex vivo* lineage specification of naïve CD4⁺ T cells toward the T_H_17 cell lineage.

**Figure 1.**
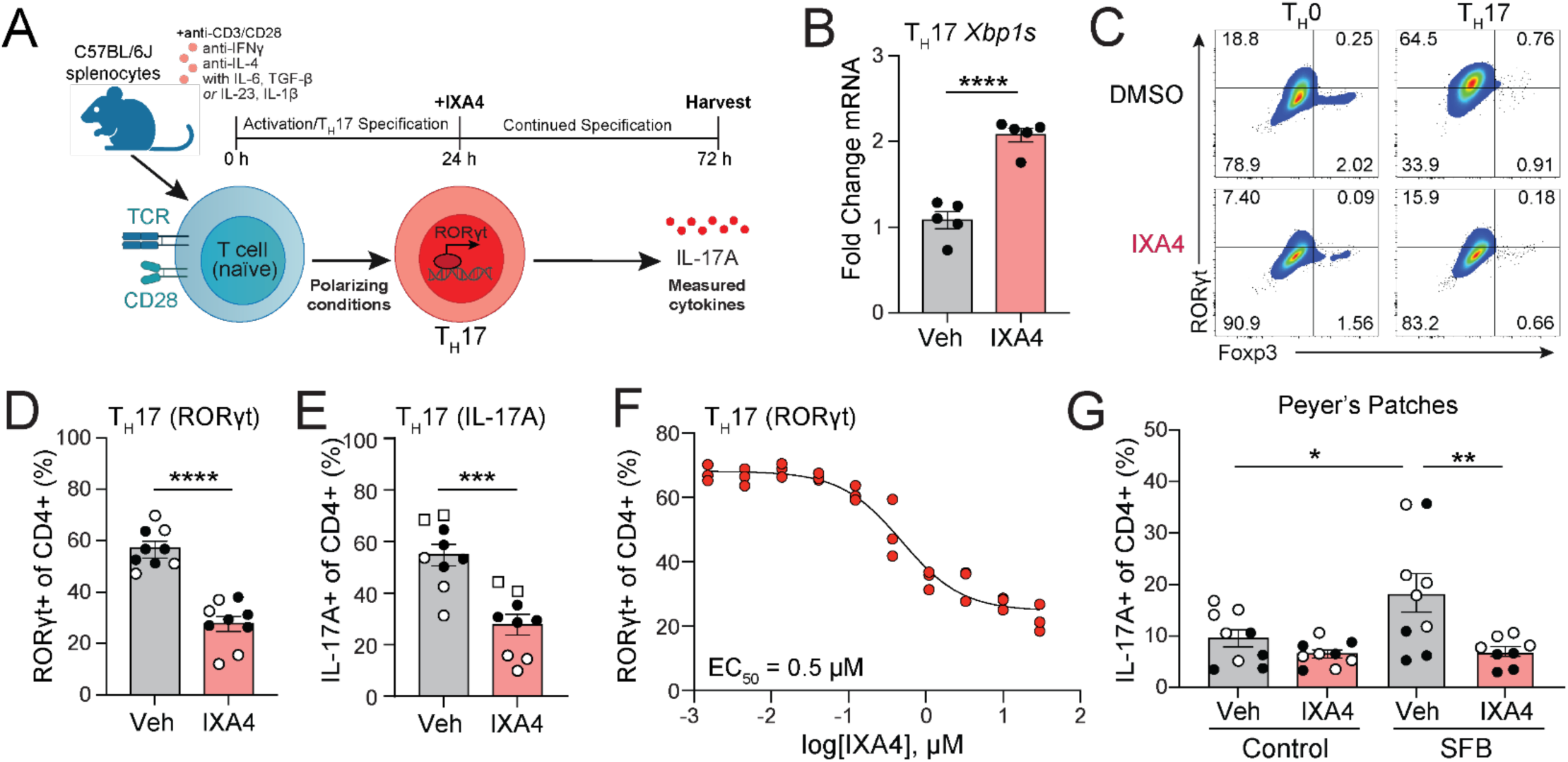
The IRE1/XBP1s activator IXA4 selectively impairs *ex vivo* and *in vivo* differentiation of CD4^+^ T_H_17 cells. **A.** Schematic of CD4^+^ T_H_17 lineage specification and experimental timeline for *ex vivo* differentiation of CD4^+^ T cells. This protocol was used for all described experiments unless otherwise indicated. **B**. Expression, measured by RT-qPCR, of *Xbp1s* transcript levels in T_H_17 cells differentiated for 24 h and then treated for 6 h with IXA4 (10 μM) or vehicle. Error bars show SEM for n=5 biological replicates. **C.** Representative flow cytometry plots of RORγt and Foxp3 staining in CD4^+^ T cells polarized under T_H_0 (polarized with IL-2) or T_H_17 conditions in the presence of IXA4 (10 μM) or vehicle. **D.** Quantification of RORγt^+^ cells in CD4^+^ T cells polarized under T_H_17 conditions in the presence of IXA4 (10 μM) or vehicle. Error bars show SEM for n=9 biological replicates across 2 independent experiments. **E**. Quantification of IL-17A^+^ CD4^+^ T cells polarized under T_H_17 conditions in the presence of IXA4 (10 µM) or vehicle. Error bars show SEM for n=9 biological replicates across 3 independent experiments. **F**. Dose-dependent reduction of RORγt^+^ cells in CD4^+^ T cells differentiating in T_H_17 conditions treated with the indicated concentration of IXA4. Error bars show SEM for n=3 biological replicates. **G**. Quantification of IL-17A^+^ CD4^+^ T cells in Peyer’s patches of SFB^neg^ mice daily administered SFB^pos^ fecal slurry, IXA4 (50 mg/kg), and/or vehicle by oral gavage for 10 days. Error bars show SEM for n=9 biological replicates across 2 independent experiments. *p<0.05, **p<0.01, ***p<0.001, ****p<0.0001 for two tailed unpaired t-test (**B, D, E**) or one-way ANOVA (**G**).

We next used segmented filamentous bacteria (SFB) colonization of mice to probe the potential for IXA4 to suppress T_H_17 cell differentiation *in vivo*. Colonization of the small intestine with the commensal bacterium SFB promotes the expansion of T_H_17 cells within intestinal lymphoid tissues such as the Peyer’s patches (PP)^20,60^. Consistent with this, we observed increased frequency of IL-17A^+^ producing CD4^+^ T cells in PP isolated from SFB-free mice dosed orally with feces from an SFB-positive mouse daily for 10 days (**Fig. S1M-N**). Daily oral gavage administration of IXA4 reduced the SFB-dependent increase of IL-17A-producing CD4^+^ T cells in the PP (**Fig. 1G, Fig. S1O**). These results indicate that IXA4 treatment can limit SFB-dependent increases of T_H_17 cells *in vivo*.

### IXA4-induced reduction in T_H_17 differentiation requires IRE1 activity

IXA4 was identified as a compound that preferentially activates the IRE1/XBP1s signaling arm of the UPR^53,54^. IRE1/XBP1s signaling is increased during normal T_H_17 differentiation, with the highest levels of signaling observed ∼20 h after the initiation of differentiation (**Fig. S2A-B**)^61^. The inhibition of T_H_17 differentiation by IXA4 suggests that the compound either further boosts IRE1 activity already present in differentiating cells or acts through an alternative mechanism or target. Thus, we sought to determine whether the effect on T_H_17 differentiation afforded by IXA4 could be attributed to increased IRE1 signaling. RNAseq transcriptional profiling showed that treatment of differentiating T_H_17 cells for 6 h with IXA4 increased UPR signaling (**Fig. 2A**, **Table S1**). We observed increased expression of a gene set comprising IRE1/XBP1s target genes, without observing significant increases in gene sets regulated by the PERK/ATF4 signaling arm of the UPR (**Fig. 2B-C**, **Fig. S2-D**, **Table S1**)^45,62–65^. Co-treatment with the IRE1/RNase inhibitor STF-080310 blocked the IXA4-induced increases in the expression of IRE1/XBP1s target genes, indicating that this increase can be attributed to IRE1 activity (**Fig. 2B-C**, **Fig. S2C**)^66^. In contrast, PERK/ATF4 target gene expression was increased in cells co-treated with IXA4 and STF-080310 (**Fig. 2B**, **Fig. S2D**). Collectively, these results show that IXA4 retains the ability to preferentially activate the IRE1/XBP1s arm of the UPR in differentiating T_H_17 cells.

**Figure 2.**
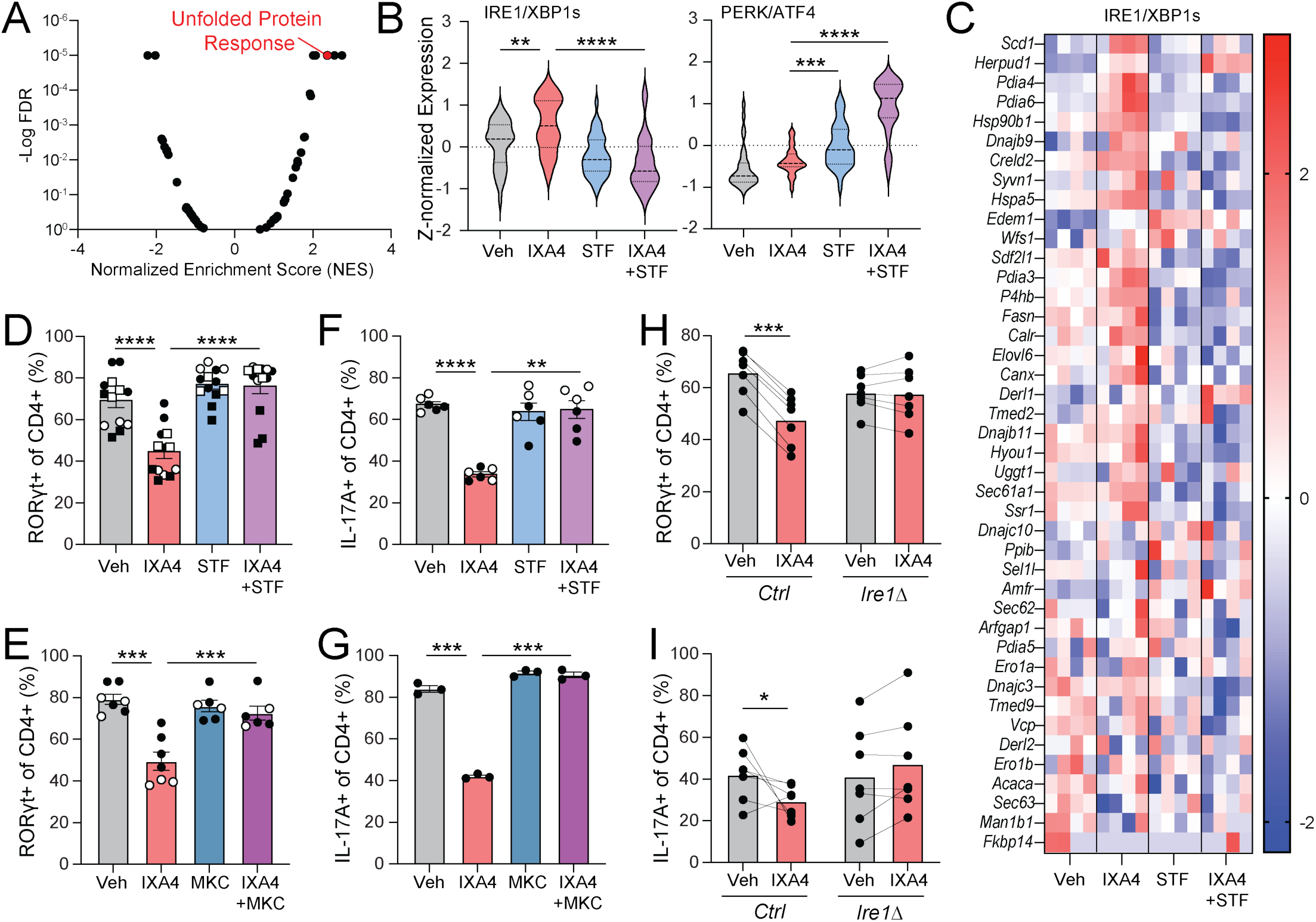
IXA4-mediated reductions in T_H_17 differentiation require IRE1 activity. **A.** Gene set enrichment analysis (GSEA) of RNAseq from CD4^+^ T cells differentiated for 24 h in T_H_17 polarizing conditions and then treated for an additional 6 h with vehicle or IXA4 (10 µM). The full GSEA can be found in **Table S1**. **B**. Z-normalized expression of gene sets comprising IRE1/XBP1s target genes^45,62–65^ or PERK/ATF4 target genes^45^ in CD4^+^ T cells differentiated for 24 h and then treated for an additional 6 h with vehicle or IXA4 (10 µM). Full gene sets are shown in **Table S1**. **C.** Heat map of Z-normalized gene expression of IRE1/XBP1s target gene sets described in panel (**B**). **D-G.** Percent RORγt^+^ (**D,E**) or IL-17A^+^ (**F,G**) CD4^+^ T cells differentiated in T_H_17 polarizing conditions in the presence of IXA4 (10 µM), STF-080310 (64 µM), or MKC-3946 (10 µM). Error bars show SEM for n= 3-14 individual replicates across 1-4 individual experiments. **H,I.** Percent RORγt^+^ (**H**) or IL-17A^+^ (**I**) in *Ctrl* or *Ire1Δ* CD4^+^ T cells differentiated in T_H_17 polarizing conditions and treated with IXA4 (10 µM) or vehicle, as indicated. Error bars show SEM for n=7 replicates across three individual experiments. *p<0.05, **p<0.01, ***p<0.005, ****p<0001 for one-way ANOVA (panels **B, D, E, F, G**) or paired t-test (panel **H,I**).

We next defined the dependence of IXA4-induced reductions in T_H_17 differentiation on IRE1 activity using three structurally distinct IRE1 RNase inhibitors: 4µ8c, STF-080310, and MKC-3946^66–69^. Treatment with 4µ8c reduced viability and decreased RORγt expression in differentiating T_H_17 cells (**Fig. S2E-F**). This is consistent with previous work showing that treatment with this compound blocked the *ex vivo* differentiation of T_H_17 cells^52^. In contrast, treatment with STF-080310 or MKC-3946 alone did not influence the viability or RORγt expression in differentiating T_H_17 cells (**Fig. S2E-F**). However, co-treatment with STF-080310 or MKC-3946 inhibited IXA4-induced increases in *Xbp1s* in differentiating T_H_17 cells (**Fig. S2G**). Thus, we used these two IRE1 RNase inhibitors to probe the importance of IRE1 activity in IXA4-dependent reductions in T_H_17 differentiation. Co-treatment with either STF-080310 or MKC-3946 prevented IXA4-dependent reductions in RORγt expression and IL-17A production in differentiating T_H_17 cells (**Fig. 2D-G**). This indicates that IRE1 activity is required for the IXA4-dependent reductions in differentiation of this T cell subset.

To further test the dependence of IXA4-dependent reductions in T_H_17 differentiation on IRE1 activity, we isolated naïve CD4^+^ T cells deficient in *Ern1* (the gene encoding IRE1) from the spleens of *Rosa26^CreERT^*^2^ *Ern1^fl/fl^* mice (referred to as *Ire1Δ*) and *Rosa26^CreERT2^ Ern1^fl/wt^* littermate controls (*Ctrl*) one week after tamoxifen-induced Cre activation to avoid confounding effects from chronic loss of IRE1. We confirmed *Ern1* deletion was successful as the IXA4-induced increase in *Xbp1s* was blocked in *Ire1Δ* cells, as compared to *Ctrl* cells (**Fig. S2H**). Importantly, *Ire1Δ* cells showed no change in viability or activation status relative to *Ctrl* cells in T_H_17 polarizing conditions (**Fig. S2I-J**). IXA4 treatment did not reduce RORγt expression or IL-17A production in differentiating *Ire1Δ* T_H_17 cells, as compared to *Ctrl* cells, further confirming that IXA4-induced suppression of T_H_17 differentiation requires IRE1 activity (**Fig. 2H-I**).

### IXA4-induced IRE1 activation decreases intracellular cholesterol in differentiating T_H_17 cells

To define the mechanism of action whereby IXA4-induced IRE1 activation blocks T_H_17 cell differentiation without the confounding secondary effects that may accumulate as differentiation proceeds, we focused on early responses to the compound treatment. Treatment with IXA4 for 24 h was sufficient to reduce expression of RORγt under T_H_17 polarizing conditions (**Fig. S3A**). Thus, we performed RNAseq on differentiating T_H_17 cells treated for 24 h with IXA4 in the presence or absence of the IRE1 RNase inhibitor STF-083010 to further probe IXA4-dependent gene expression changes in these cells (**Table S2**). Gene set enrichment analysis (GSEA) showed that IXA4 treatment for 24 h increased UPR signaling (**Fig. 3A**, **Table S2**). Treatment with STF-080310 reduced expression of IRE1/XBP1s target genes both in the presence and absence of IXA4 (**Fig. S3B-C**, **Table S2**), reflecting the basal activation of this pathway during T_H_17 cell differentiation (**Fig. S2A-B**). Intriguingly, unlike what we observed at 6 h, IXA4 treatment for 24 h increased expression of PERK/ATF4 target genes in differentiating T_H_17 cells through a mechanism refractory to co-treatment with STF-080310 (**Fig. S3B-C**). Similar results were observed in cells co-treated with IXA4 and the alternative IRE1 inhibitor MKC-3946 (**Fig. S3D-E, Table S3**). Co-treatment with either the PERK kinase inhibitor AMG44 or the PERK signaling inhibitor ISRIB blocked IXA4-dependent increases in the expression of PERK/ATF4 target genes, confirming that this increase can be attributed to PERK activation (**Fig. S3F-G**)^69–71^. These results indicate that chronic treatment with IXA4 can also lead to increased PERK/ATF4 signaling in differentiating T_H_17 cells.

**Figure 3.**
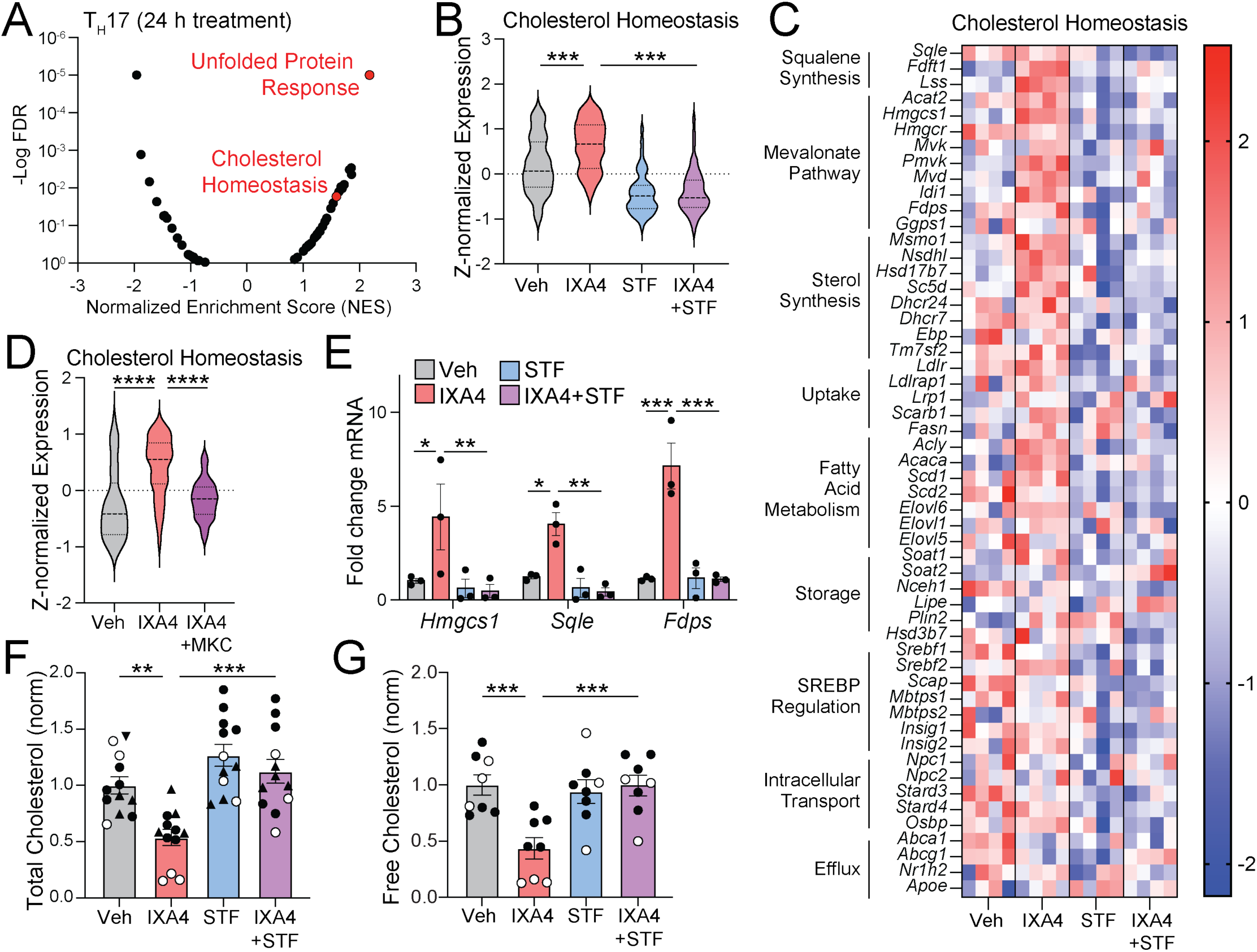
IXA4-mediated IRE1 activation depletes intracellular cholesterol in differentiating T_H_17 cells. **A.** GSEA of hallmark gene sets in CD4^+^ T cells polarized under T_H_17 conditions for 24 h and treated for 24 h with IXA4 (10 µM) or vehicle. Full GSEA is shown in **Table S2**. **B.** Z-normalized expression of the cholesterol homeostasis gene set in differentiating T_H_17 cells treated for 24 h with IXA4 (10 µM) and/or STF-083010 (64 µM). The full gene set is shown in **Table S2**. **C.** Heatmap showing Z-normalized expression of individual cholesterol homeostasis genes, measured by RNAseq, across the conditions described in (**B**). Each column represents an individual biological replicate. **D.** Z-normalized expression of the cholesterol homeostasis gene set in differentiating T_H_17 cells treated for 24 h with IXA4 (10 µM) and/or MKC-3946 (10 µM). The full gene set is shown in **Table S3**. **E.** Expression of the cholesterol biosynthesis genes *Hmgcs1, Sqle, and Fdps,* measured by RT-qPCR, in differentiating T_H_17 cells treated for 48 h with IXA4 (10 µM) and/or STF-083010 (64 µM). Error bars show SEM for n=3 biological replicates. **F,G.** Normalized total (**F**) and free (**G**) cholesterol in differentiating T_H_17 cells treated for 48 h with IXA4 (10 µM) and/or STF-083010 (64 µM). Error bars show SEM for n=8 or 12 biological replicates across 2 or 3 individual experiments. *p < 0.05, **p < 0.01, ***p < 0.001, ****p < 0.0001 for one-way ANOVA (**B, D, F, G**) or two-way ANOVA (**E**).

Notably, treatment of differentiating T_H_17 cells with IXA4 for 24 h increased expression of a gene set comprising cholesterol homeostasis genes (**Fig. 3A-D**, **Tables S2-S3**). Cholesterol metabolism is a central regulator of T_H_17 cell biology^34,72^. Differentiation induces cholesterol biosynthesis, and sterol pathway intermediates can serve as endogenous RORγt agonists, directly linking metabolic flux to this lineage-defining transcription factor^33,34,73^. Sterol-sensing pathways, including those regulated by LXR and the SREBPs, regulate T_H_17 differentiation and effector programs, and cholesterol and its precursors support the membrane composition, signaling capacity, and biosynthetic demands of rapidly proliferating effector T cells^74,75^. We found that IXA4-mediated increases in cholesterol homeostasis gene expression were blocked by co-treatment with the IRE1 RNase inhibitor STF-080310, indicating that this effect requires IRE1 activity (**Fig. 3B-C**, **Table S2**). Similar results were observed in differentiating cells treated for 24 h with IXA4 in the absence or presence of the alternative IRE1 RNase inhibitor MKC-3946 (**Fig. 3D**, **Table S3**). IXA4 treatment primarily increased expression of cholesterol biosynthesis genes such as *Hmgcs1, Sqle*, and *Fdps* (**Fig. 3C-E**, **Fig. S3H**). Cholesterol biosynthesis is tightly regulated, with sterol-sensing pathways inducing cholesterol biosynthesis genes in response to deficits in intracellular cholesterol^76,77^. Therefore, the induction of these genes by IXA4 suggested that IXA4 treatment reduced intracellular cholesterol^78^. To test this, we measured total and free intracellular cholesterol in differentiating T_H_17 cells treated with IXA4 or vehicle. IXA4 treatment reduced total and free cholesterol in differentiating T_H_17 cells in an IRE1-dependent manner, as co-treatment with the IRE1 RNase inhibitor STF-080310 blocked IXA4-induced reductions in cholesterol (**Fig. 3F-G**). As T_H_17 cells are exquisitely sensitive to cholesterol availability, this decrease suggests a mechanism to explain the impaired differentiation of this T cell subset afforded by IXA4 treatment.

### IXA4 suppresses T_H_17 differentiation by limiting cholesterol biosynthesis

3-hydroxy-3-methylglutaryl-CoA reductase (HMGCR) is the rate-limiting enzyme in cholesterol synthesis. HMGCR is an ER membrane protein that can be targeted for proteasomal degradation in response to changes in intracellular cholesterol through ER-associated degradation (ERAD)^79–81^, a process whose capacity is regulated by the IRE1/XBP1s signaling pathway^65,82,83^. Therefore, we hypothesized that the reduction in intracellular cholesterol observed in differentiating T_H_17 cells treated with IXA4 could result from increased proteasome-mediated degradation of HMGCR. Indeed, we found that IXA4 treatment for 24 h reduced HMGCR protein levels in differentiating T_H_17 cells, as measured by both immunoblotting and flow cytometry (**Fig. 4A-B**). Co-treatment with the IRE1 inhibitor STF-080310 blocked IXA4-induced reductions of HMGCR (**Fig. 4B**). Similarly, co-treatment with either of the two proteasome inhibitors bortezomib (BTZ) or MG132 for 4 h prevented IXA4-mediated reductions in HMGCR, consistent with HMGCR reduction being the result of its enhanced proteasomal degradation (**Fig. 4C-D; Fig.S4A**). Consistent with a requirement for proteasomal degradation in this phenotype, co-treatment with either bortezomib or MG132 also prevented IXA4-dependent reductions in RORγt expression in differentiating T_H_17 cells (**Fig. 4F-G**). These results indicate that IXA4-induced IRE1 activation stimulates proteasomal degradation of HMGCR, the rate-limiting cholesterol biosynthesis enzyme.

**Figure 4.**
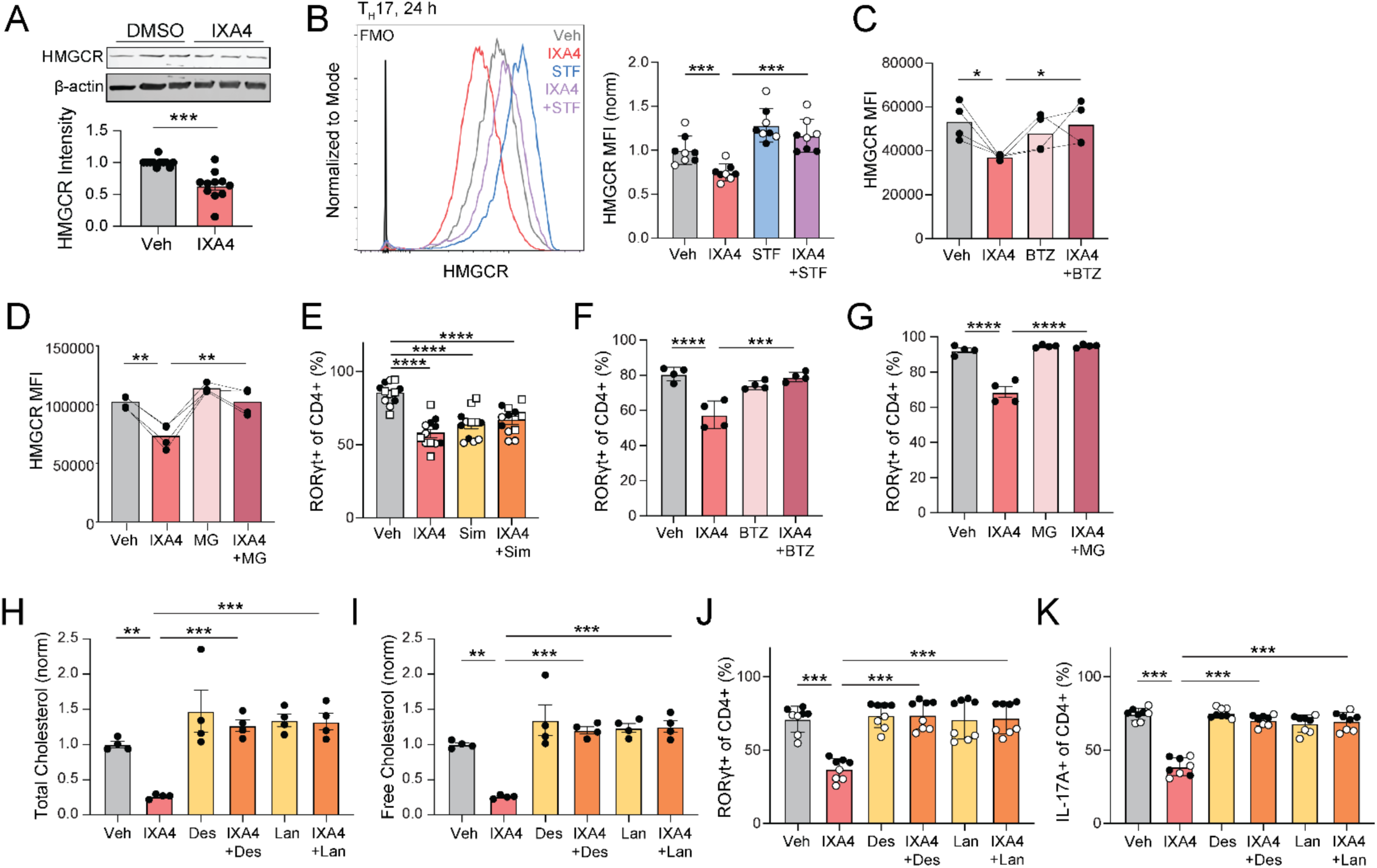
Co-treatment with cholesterol precursors prevents IXA4-mediated reductions in T_H_17 differentiation. **A.** Representative immunoblot and quantification of HMGCR in T_H_17 cells treated for 24 h with IXA4 (10 µM) or vehicle. Error bars show SEM for n=12 replicates across 4 individual experiments. **B.** Representative histograms and quantification of mean fluorescence intensity (MFI) for HMGCR, measured by flow cytometry, in differentiating T_H_17 cells treated for 24 h with IXA4 (10 µM) in the absence or presence of STF-083010 (64 µM). Error bars show SEM for n=8 replicates across 2 independent experiments. **C,D.** HMGCR levels, measured by flow cytometry, in differentiating T_H_17 cells treated with IXA4 (10 µM) for 24 h and then treated an additional 4 h with bortezomib (1 nM; **C**) or MG132 (10 µM; **D**). Error bars show SEM for n= 4 biological replicates. **E.** Percent RORγt^+^ cells in differentiating T_H_17 cells treated with IXA4 (10 µM) and/or simvastatin (1 µM). Error bars show SEM for n=12 replicates across 3 independent experiments. **F,G.** Percent RORγt^+^ cells in differentiating T_H_17 cells treated with IXA4 (10 µM) and then treated for an additional 4 h with bortezomib (**F**) or MG132 (**G**). Error bars show SEM for n=4 replicates. **H,I.** Normalized total (**H**) and free (**I**) cholesterol in differentiating T_H_17 cells treated with IXA4 (10 µM) in the absence or presence of desmosterol (Des, 10 µM) or lanosterol (Lan, 10 µM). Error bars show SEM for n=4 replicates. **J, K.** Percent RORγt^+^ (**J**) and IL-17A^+^ (**K**) cells in differentiating T_H_17 cells treated with IXA4 (10 µM) in the absence or presence of desmosterol (10 µM) or lanosterol (10 µM). Error bars show SEM for n=8 replicates across 2 independent experiments. *p < 0.05, **p < 0.01, ***p < 0.001, ****p < 0.0001 for unpaired t-test (**A**) or ANOVA (**B-K**).

If reduced HMGCR levels underlie the effects of IXA4, then pharmacologic inhibition of HMGCR activity should have similar effects on T_H_17 differentiation. Indeed, statin-induced inhibition of HMGCR suppresses differentiation of T_H_17 cells^84–86^. Consistent with this, treatment with simvastatin decreased RORγt expression in differentiating T_H_17 cells comparably to IXA4 treatment, although we did observe reductions in viability of simvastatin-treated cells (**Fig. 4E**, **Fig. S4B-C**). Importantly, co-treatment with IXA4 and simvastatin did not show further reductions in RORγt expression (**Fig. 4E**). This lack of additivity is consistent with an epistatic relationship where IXA4 and simvastatin both suppress T_H_17 differentiation by inhibiting HMGCR activity, rather than through independent mechanisms.

The increased degradation of HMGCR in cells treated with IXA4 supports a model whereby increased IRE1 activation reduces cholesterol biosynthesis below a level required to support T_H_17 cell differentiation. If reduced cholesterol biosynthesis is indeed the mechanism by which IXA4 suppresses T_H_17 differentiation, then supplying sterol intermediates generated downstream of HMGCR should rescue differentiation in IXA4-treated cells. To test this, we co-treated differentiating T_H_17 cells with IXA4 and the cholesterol precursors desmosterol or lanosterol, two sterols generated downstream of HMGCR in the cholesterol biosynthesis pathway. Co-treatment with these cholesterol precursors did not influence IXA4-dependent increases in *Xbp1s* or the induction of IRE1/XBP1s target genes (**Fig. S4D-E, Table S4**). However, co-treatment of differentiating T_H_17 cells with IXA4 and either desmosterol or lanosterol restored intracellular cholesterol levels and blocked IXA4-induced expression of cholesterol biosynthesis genes such as *Sqle* or *Msmo1* (**Fig. 4H-I; Fig. S4F**). Moreover, co-treatment of differentiating T_H_17 cells with desmosterol or lanosterol blocked IXA4-induced reductions in RORγt expression and IL-17A production in differentiating T_H_17 cells (**Fig. 4J-K**). These results indicate that exogenous addition of cholesterol precursors generated downstream of HMGCR in the cholesterol biosynthesis pathway restores differentiation of IXA4-treated T_H_17 cells.

### The alternative IRE1/XBP1s activator IXA62 also decreases T_H_17 differentiation by rewiring cholesterol metabolism

Apart from IXA4, we previously identified the compound IXA62 as a structurally distinct compound that also selectively activates the IRE1/XBP1s signaling arm of the UPR (**Fig. S5A**)^87^. We sought to determine if IXA62-mediated IRE1 activation, like IXA4, reduced T_H_17 differentiation by remodeling cholesterol metabolism. We confirmed that IXA62 treatment for 6 h preferentially increased expression of IRE1/XBP1s target genes in differentiating T_H_17 cells (**Fig. 5A-C, Table S5**). Further, we found that treatment with IXA62 reduced expression of RORγt and IL-17A production in differentiating T_H_17 cells to levels similar to those observed in IXA4-treated cells (**Fig. 5D-E**). Co-treatment with the IRE1 RNase inhibitors STF-080310 or MKC-3946 blocked this reduction (**Fig. S5B-D**). In addition, IXA62 treatment did not influence RORγt expression or IL-17A production in *Ern1*-deficient CD4^+^ T cells incubated in T_H_17 polarizing conditions compared to littermate controls (**Fig. 5F-G**). These results demonstrate that, like IXA4, IXA62 treatment reduces differentiation of T_H_17 cells through an IRE1-dependent mechanism.

**Figure 5.**
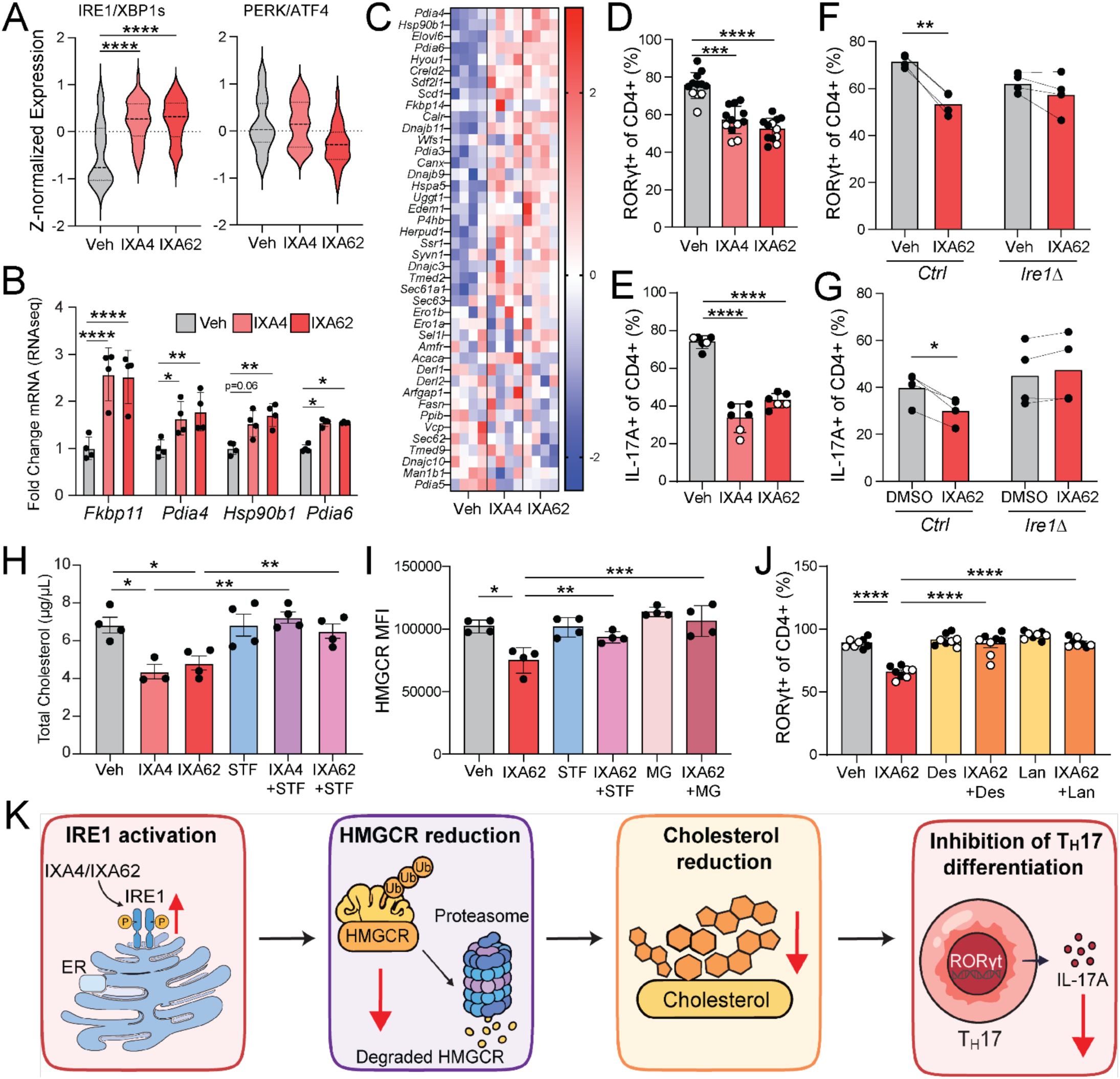
IXA62-mediated IRE1 activation reduces T_H_17 differentiation by rewiring cholesterol metabolism. **A.** Z-normalized expression of gene sets comprising IRE1/XBP1s target genes (left) and PERK/ATF4 target genes (right) in CD4⁺ T cells polarized under T_H_17 conditions and treated for 6 h with IXA4 (10 µM), IXA62 (10 µM), or vehicle. Full gene sets are described in **Table S5**. **B**. Expression of IRE1/XBP1s target genes, measured by RNAseq, in conditions described in (**A**). **C**. Heat map of IRE1/XBP1s target genes across the conditions described in (**A**). Each column represents an individual biological replicate. **D,E.** Percent RORγt⁺ (**D**) and IL-17A⁺ (**E**) cells in CD4⁺ T cells polarized under T_H_17 conditions and treated with IXA4 (10 µM), IXA62 (10 µM), or vehicle, as indicated. Error bars show SEM for n=6-12 replicates across 2 independent experiments. **F,G.** Percent RORγt⁺ (**F**) and IL-17A⁺ (**G**) cells in CD4⁺ T cells isolated from *Ctrl* or *Ire1Δ* mice, polarized under T_H_17 conditions and treated with IXA62 (10 µM) or vehicle. Connecting lines indicate paired samples from the same experiment. Error bars show SEM for n=3 replicates. **H.** Total cholesterol in differentiating T_H_17 cells treated with IXA4 (10 µM) or IXA62 (10 µM) in the absence or presence of STF-083010 (64 µM). Error bars show SEM for n=4 biological replicates. **I.** HMGCR levels, measured by flow cytometry, in differentiating T_H_17 cells treated with IXA62 (10 µM), STF-083010 (64 µM), or MG132 (10 µM). STF-080310 was administered at the same time as IXA62, while MG132 was added for the final 4 h of treatment. Error bars show SEM for n=4 replicates. **J.** Percent RORγt⁺ cells in differentiating T_H_17 cells treated with IXA62 (10 µM) in the absence or presence of desmosterol (Des, 10 µM) or lanosterol (Lan, 10 µM). Error bars show SEM for n=8 replicates across 2 independent experiments. **K.** Schematic for the proposed model of IXA4-mediated reductions in T_H_17 differentiation. \**p* < 0.05, \*\**p* < 0.01, *** p < 0.001, \*\*\*\**p* < 0.0001 for one-way ANOVA (**B, D, E, H-J**) or paired t-test (**F, G**).

IXA62 treatment also reduced cholesterol levels in differentiating T_H_17 cells to levels similar to those observed with IXA4 (**Fig. 5H**). Co-treatment with the IRE1 RNase inhibitor STF-080310 blocked this reduction, confirming that this effect is attributed to IRE1 activity. Further, we found that IXA62 decreased HMGCR in differentiating T_H_17 cells through a process sensitive to co-treatment with the IRE1 RNase inhibitor STF-080310 or the proteasome inhibitor MG132 (**Fig. 5I**, **Fig. S5E**). Finally, we show that co-treatment with the cholesterol precursors desmosterol or lanosterol rescued both IXA62-dependent reductions in cholesterol and expression of RORγt in differentiating T_H_17 cells (**Fig. 5J**, **Fig. S5F-G**). Collectively, these results demonstrate that pharmacologic IRE1 activation with the structurally distinct compound IXA62 reduces T_H_17 differentiation by promoting HMGCR proteasomal degradation and reducing intracellular cholesterol.

## DISCUSSION

Here, we show that pharmacologic IRE1 activation preferentially suppresses the differentiation of CD4^+^ T_H_17 cells (**Fig. 5K**). We demonstrate that this suppression is mediated through a process involving the increased proteasomal degradation of the rate-limiting cholesterol biosynthesis enzyme HMGCR, thereby reducing cholesterol biosynthesis in differentiating T_H_17 cells. Since T_H_17 differentiation is highly dependent on cholesterol biosynthesis for its specification, this reduction inhibits the differentiation of T_H_17 cells without impacting differentiation of other CD4^+^ T cell subsets. These results identify IRE1 activation as a previously unanticipated metabolic checkpoint in T_H_17 differentiation that can be pharmacologically accessed to limit the differentiation of this pro-inflammatory T cell subset.

The sensitivity of T_H_17 cells to pharmacologic IRE1 activation is intriguing given that IRE1 pathway activity increases during normal T_H_17 cell differentiation^61^. Nonetheless, we show that neither pharmacologic nor genetic inhibition of IRE1 activity affects T_H_17 cell differentiation, suggesting that endogenous IRE1 activity is dispensable for this process. Consistent with our findings, previous results show that mice lacking *Ern1* (*Ire1α*) in RORγt-expressing cells showed no change in the RORγt^+^ T cell pool in the PP, despite deletion occurring early during T cell development, indicating that IRE1 is also dispensable for the generation or maintenance of intestinal RORγt^+^ T cells *in vivo*^88,89^. These findings contrast with a previous study reporting reduced T_H_17 responses in a fungal asthma model as well as impaired T_H_17 differentiation, following genetic or pharmacologic IRE1 inhibition^52^. Whether these discrepancies reflect context-dependent requirements for IRE1 signaling or developmental effects arising from early deletion of *Ern1 (Ire1α)* in T cells remains unclear. Importantly, this last study^52^ relied on the *in vitro* use of IRE1 RNase inhibitor 4µ8c, which we show has effects on CD4^+^ T cell viability and T_H_17 differentiation that are not observed with two structurally distinct IRE1 RNase inhibitors, STF-080310 and MKC-3946. These differences underscore the importance of using multiple different approaches to establish the significance of IRE1 signaling in differentiating cells to minimize the potential contributions of off-target activities that could confound observed results. Our results indicate that IRE1 activity is not essential for T_H_17 differentiation but instead highlight how enhancing the activity of this UPR pathway can be used to preferentially reduce differentiation of this pro-inflammatory T cell subset.

We show that pharmacologic IRE1 activation suppresses T_H_17 cell differentiation through proteasomal degradation of HMGCR, the rate-limiting enzyme in cholesterol biosynthesis, and subsequent reductions in intracellular cholesterol. HMGCR is a known substrate of ERAD, a process governed largely by IRE1/XBP1s-dependent transcriptional regulation of ERAD components including the E3 ligase HRD1, which has previously been implicated in HMGCR degradation^81,90^. Canonically, HMGCR degradation is normally induced by elevated intracellular sterols, serving as a negative feedback regulator that restrains cholesterol biosynthesis when levels are high. Given that IXA4 treatment induces the transcriptional upregulation of IRE1/XBP1s genes, it is possible that this response also generates signals that promote HMGCR degradation through ERAD, ultimately resulting in reduced intracellular cholesterol levels. This suggests a previously unanticipated mode of HMGCR regulation that can be accessed through IRE1 activation in differentiating T_H_17 cells to control cholesterol biosynthesis and subsequently the differentiation of this T cell lineage.

We show that IXA4 selectively suppresses T_H_17 cell differentiation, with no substantial effect on the differentiation of other CD4^+^ T cell subsets *in vitro*. This selectivity may reflect a particular dependence of T_H_17 cells on cholesterol biosynthesis, which is upregulated during differentiation and generates sterol intermediates that support T_H_17 lineage commitment^33,34^. Consistent with this, genetic or pharmacologic disruption of cholesterol biosynthetic enzymes, including SC4MOL, impairs T_H_17 differentiation without affecting T_H_1 or T_reg_ cell differentiation^33,34^. Similarly, statins have been shown to suppress T_H_17 cell differentiation while favoring T_reg_ cell differentiation^85,91^. Together, these findings suggest that the selective sensitivity of T_H_17 cells to IRE1 activation may arise from distinct metabolic requirements during differentiation. The ability of IXA4 to preferentially suppress T_H_17 differentiation can be attributed to our addition of our pharmacologic IRE1 activators 24 h after T cell activation, providing an opportunity to specifically access the preferential sensitivity of T_H_17 cells to reductions in cholesterol. This indicates a unique role for IRE1 activation as a metabolic checkpoint in T_H_17 differentiation that could be selectively accessed to limit the differentiation of the CD4^+^ T cell subset independent of other lineages.

The aberrant proliferation of T_H_17 cells is implicated in the onset and pathogenesis of many inflammatory diseases, including psoriasis, multiple sclerosis, and inflammatory bowel disease^22,23^. Our results highlight an opportunity to limit the pathologic accumulation of pro-inflammatory T_H_17 cells in these diseases using pharmacologic IRE1 activators. We demonstrate this potential by showing that treatment with the IRE1 activator IXA4 decreases T_H_17 cell accumulation in Peyer’s patches from mice colonized with SFB. An advantage of this approach is that, apart from limiting pro-inflammatory T_H_17 cells, pharmacologic IRE1 activation can also promote adaptive remodeling of multiple other cell types and tissues that can integrate to broadly mitigate pathologies linked to these pro-inflammatory diseases. This type of multi-tissue benefit has been previously demonstrated in models of obesity-diabetes where IXA4-induced IRE1 activation promotes adaptive remodeling of both liver and pancreas that integrate to improve multiple metabolic parameters and overall organismal health^54^. Our results further expand the therapeutic potential for pharmacologic IRE1 activation by showing their potential for directly limiting T_H_17 cell differentiation, revealing new ways in which the systemic administration of these compounds can improve overall organismal health in the context of disease. As we continue developing this class of compounds, we will expand these studies to probe their potential for multi-tissue benefits in inflammatory diseases, further highlighting the therapeutic benefits for systemic IRE1 activation in human disease.

## METHODS

### Mice

C57BL/6 mice were bred in-house by the Rodent Breeding Colony (RBC) at Scripps Research. C57BL/6NTac were purchased from Taconic Biosciences for SFB infection model experiments. C57BL/6J-*Ptprc^em6Lutzy^*/J (JAXBoy, Jackson Laboratory, #033076) mice were obtained and crossed with B6.129-*Gt(ROSA)26Sor^tm1(cre/ERT2)Tyj^*/J (R26-CreERT2, Jackson Laboratory, #008463). The resulting offspring were further crossed with *Ern1^fl/fl^*mice^92^ to generate an inducible deletion of *Ern1*. All animal experiments were performed in accordance with protocols approved by the Institutional Animal Care and Use Committee (IACUC) and complied with Scripps Research institutional guidelines.

### Chemicals and Reagents

The IRE1/XBP1s activators described herein were purchased: IXA62 (Enamine, Cat #T5347952). Larger quantities of IXA4 required for *in vivo* studies were synthesized by Otava. We confirmed the identity of all compounds by NMR, HPLC, and mass spectrometry. 4μ8c (EMD, Cat #412512), STF-083010 (Selleck chemicals, Cat #S7771), MKC3946 (Fisher Scientific, Cat #HY-19710), desmosterol (Sigma, Cat # D6513), lanosterol (Avanti Research, Cat #700099P), bortezomib (Fisher, Cat #50-431-40001), MG132 (Selleck chemicals, Cat #S2619), ISRIB (Sigma, Cat #SML0843), AMG44 (Selleck chemicals, Cat #E1761), Simvastatin (Fisher, Cat #AC458840010) were purchased from commercial vendors.

### T cell differentiation assays

Naïve CD4^+^ cells were isolated from splenocytes using MojoSort Mouse Naïve CD4 T Cell Isolation Kit (BioLegend, Cat #480040). Isolated T cells were activated with anti-CD3 (0.25 µg/mL, BioXCell, Cat #BE0001-2) and anti-CD28 (1 µg/mL, BioXCell, Cat #BE0015-1) and differentiated into T_H_1 cells with IL-12 (10 ng/mL, PeproTech, Cat #210-12P80H), IL-2 (50 µg/mL, PeproTech, Cat #200-02) and anti-IL-4 (2 µg/mL, BioXCell, Cat #BE0045). T_H_2 cells were differentiated with IL-2 (50 µg/mL), IL-4 (2 ng/mL, PeproTech, Cat #214-14), and anti-IFNγ (2 µg/mL, BioXCell, Cat #BE0055). T_H_17 cells were differentiated with anti-IL-4 (2 µg/mL), and anti-IFNγ (2 µg/mL) and TGF-β (0.3 ng/mL, PeproTech, Cat #100-21C) and IL-6 (10 ng/mL, PeproTech, Cat #216-16), or IL-1β (20 ng/mL, PeproTech, Cat #211-11B), and IL-23 (25 ng/mL, eBioscience, Cat #14-8231-63). T_reg_ cells were differentiated with IL-2 (50 µg/mL), TGF-β (5 ng/mL), anti-IL-4 (2 µg/mL), and anti-IFNγ (2 µg/mL) for 3 days.

For experiments involving compound treatments, activators and inhibitors (or sterols) were added 24 hours after initial T cell activation. Cells were harvested on day 2 or day 3 for mRNA and cytokine analysis. Cells were fixed and permeabilized for transcription factor staining using eBioscience Foxp3 / Transcription Factor Staining Buffer Set (Thermo Scientific). T cell cytokine production was assessed by flow cytometry after 3 h of stimulation with PMA (100 ng/mL), ionomycin (1 µg/mL), brefeldin A (2 µg/mL), and monensin (2 µM). Intracellular cytokine staining was done using labeled antibodies after fixation and permeabilization (CytoFix, BD Biosciences). Data was obtained using a Cytek Aurora spectral flow cytometer and analyzed by FlowJo v. 10 software.

### Flow cytometry antibodies

The following antibodies were used for flow cytometry experiments: CD4 (GK1.5), CD44 (IM7), CD45 (30-F11), Foxp3 (FJK-16s), CD8α (53-6.7), IFNγ (XMG1.2), IL-17A (eBio17B7), TCRβ (H57-597), CD69 (H1.2F3), T-bet (4B10), RORγt (Q31-378), GATA-3 (TWAJ), XBP1s (143F), TCRγδ (GL-3), CD90.2 (53-2.1), CD11b (M1/70), CD62L (MEL-14), IL-4 (11B11), IL-22 (1H8PWSR), IL-13 (eBio13A), IL-10 (JES5-16E3), IL-2 (JES6-5H4), TNF (MP6-XT22), IL-5 (TRFK5), CD3ε (17A2), Zombie NIR, AlexaFluor594 Rabbit IgG (Life Technologies).

### Western Blots

Protein expression was analyzed by immunoblotting using standard protocols. Briefly, cells were lysed in ice-cold RIPA buffer supplemented with protease and phosphatase inhibitors and benzonase. Lysates were clarified by centrifugation at 16,000g for 10 min at 4 °C. Protein concentrations were determined by Bradford assay, and 30 µg of protein per sample were resolved by SDS–PAGE on polyacrylamide gels of appropriate percentage (10% or gradient). Proteins were transferred onto nitrocellulose membranes using a wet transfer system, followed by blocking in 5% (w/v) nonfat dry milk in Tris-buffered saline containing 0.1% Tween-20 (TBST). Membranes were incubated with primary antibodies diluted in blocking buffer overnight at 4 °C, washed extensively with TBST, and incubated with species-appropriate secondary antibodies for 1 h at room temperature. Immunoreactive bands were visualized using enhanced chemiluminescence and imaged on a digital imaging system. Band intensities were quantified by densitometry using ImageJ software and normalized to the indicated loading controls. Primary antibodies used: Rabbit HMGCR (Invitrogen, Cat #PA5-95846), Rabbit β-Actin (Cell Signaling, Cat #8457). Secondary antibodies used: IRDye 800CW Goat Anti-Rabbit (LICOR Biotech, Cat #926-32211).

### Sterol content assay

Cells were washed three times with PBS and lysed with organic solvents (7:11:0.1, chloroform/isopropanol/NP-40). Free (3-OH) and total (3-OH and esters) sterol levels were measured using Cholesterol/Cholesteryl Ester Assay Kit (Abcam, Cat #65359) and normalized to cholesterol standard concentrations.

### SFB colonization and T_H_17 cell induction

Segmented filamentous bacteria (SFB) colonization was established by oral gavage of recipient mice with fecal pellets derived from SFB-colonized Taconic donor mice for 10 consecutive days. Control animals were gavaged in parallel with fecal pellets from SFB-negative donors. Where indicated, in a separate gavage, mice were treated daily by oral gavage with IXA4 (50 mg kg⁻¹) or vehicle (DMSO) control throughout the colonization period. Induction of PP T_H_17 cells was assessed 10 days after initiation of SFB colonization. Fecal samples were collected daily to monitor and confirm SFB colonization over time. At experimental endpoints, mice were euthanized, and small intestine segments and Peyer’s patches were harvested for downstream analyses.

### Fecal DNA extraction for qPCR

Bacterial genomic DNA was extracted from frozen fecal samples by phenol-chloroform extraction. One fecal pellet (∼100mg) of fecal sample was suspended in a solution containing 500 µL of DNA extraction buffer (200 mM Tris, 200 mM NaCl, 20 mM EDTA), 210 µL of 20% SDS, 500 µL of phenol:chloroform:isoamyl alcohol (P:C:I) (25:24:1), and two stainless steel beads (3.2 mm diameter, BioSpec Products). Cells were lysed by mechanical disruption with a Lysera bead mill homogenizer (Biotage) for 1 minute at 6 m/s. After lysis, samples were centrifuged at 16,000 g for 5 minutes at 4°C and the DNA-containing aqueous phase was collected and subjected to two additional P:C:I extractions. DNA was precipitated with 100 µL of 3M sodium acetate pH 5.2 and 200 µL of 100% ethanol for 10 minutes followed by centrifugation for 20 minutes at 16,000 g and 4°C. DNA pellets were resuspended in TE buffer. RNA was removed with 0.1mg/mL of RNase A (Thermo Fisher) for 5 minutes at room temperature.

qPCR was conducted on the fecal DNA to determine the amount of SFB and total bacteria present in each sample by amplifying 16S rRNA genes using the bacteria-specific primers (5’-3’): *UNI-*f: ACTCCTACGGGAGGCAGCAGT, *UNI-*r: ATTACCGCGGCTGCTGCG, *SFB-*f: TGAGCGGAGATATATGGAGC, *SFB-*r: CATGCAACTATATAGCTATATGCGG. Quantitative real-time reverse transcription PCR was performed on a QuantStudio 5 Real-Time PCR system (Thermo Fisher) using SYBR Green technology. Results were analyzed and presented as the fold change relative to the internal control primer for UNI.

### RNA extraction and reverse transcription quantitative PCR (RT-qPCR)

Total RNA was extracted from pelleted cells and snap-frozen tissues using Zymo Research Quick-RNA Miniprep Kit. Complementary DNA was generated using High-Capacity cDNA Reverse Transcription Kit (Applied Biosystems). Quantitative real-time reverse transcription PCR was performed on a QuantStudio 5 Real-Time PCR system (Thermo Fisher) using SYBR Green technology. Results were analyzed and presented as the fold change relative to the internal control primer for the housekeeping gene, *Actb.* PCR primers (5’-3’) for mouse gene sequences were as follows: *Actb*-f: CATTGCTGACAGGATGCAGAAGG, *Actb*-r: TGCTGGAAGGTGGACAGTGAGG, *Xbp1s-*f: ACGAGGTTCCAGAGGTGGAG, *Xbp1s-*r: TGTCCAGAATGCCCAAAAGG, *Hmgcs1-*f: GGAAATGCCAGACCTACAGGTG, *Hmgcs1-*r: TACTCGGAGAGCATGTCAGGCT, *Trib3*-f: CTGCGTCGCTTTGTCTTCAGCA, *Trib3*-r: CTGAGTATCTCTGGTCCCACGT, *Msmo1*-f: CTGTGCAGTCATTGAGGACACC, *Msmo1*-r: GGGTTTCCAAGGGATGTGCGTA, *Cyp11a1*-f: TGCTCAACCTGCCTCCAGACTT, *Cyp11a1*-r: ACTGGCTGAAGTCTCGCTTCTG, *Fdps*-f: GGTGGTTCAGTGTCTGCTACGA, *Fdps*-r: CGCCTCATACAGTGCTTTCACC, *Sqle*-f: TGTTGCGGATGGACTCTTCTCC, *Sqle*-r: GTTGACCAGAACAAGCTCCGCA, *Ddit3*-f: GGAGCTGGAAGCCTGGTATG, *Ddit3*-r: TGTGCGTGTGACCTCTGTTG, *Asns*-f: TTACCTGTCTCTGCCGCCAGAT, *Asns*-r: CACTGAAGGCTTCTTTGGGTCG.

### RNA-Sequencing (RNAseq)

Purified RNA (50 ng/µL) was prepared using Zymo Research Quick-RNA Miniprep Kit according to the manufacturer’s instructions and submitted to Plasmidsaurus for sequencing and analysis. Values were Z-normalized across samples prior to comparison, allowing relative changes to be assessed and compared. The complete RNAseq data are deposited in the Gene Expression Omnibus (GEO) as GSE353599.

## ACKNOWLEGDEMENTS

We thank the National Institutes of Health (AG046495 to RLW; DK137470 to RLW and ES) and the Prebys Foundation (to AM) for funding. PC was funded by the Ellen Brown Scripps Foundation and the Skaggs-Oxford program, WAD was supported by the NIH (T32 TR004396 and F32 DK148145), SK was supported by a PhRMA foundation predoctoral fellowship, JS was supported by an American Diabetes Association postdoctoral fellowship, PM was supported by an American Heart Association Predoctoral Fellowship, and SD was supported by an Irvington Postdoctoral Fellowship from the Cancer Research Institute.

## SUPPLEMENTAL TABLE LEGENDS

**Table S1. RNAseq analysis of differentiating T_H_17 cells treated for 6 h with IXA4 (10 µM) and/or STF-080310 (64 µM).** This excel file contains 4 individual spreadsheets including the normalized count matrix (cpm), Gene set Enrichment Analysis (GSEA) used in **Fig. 2A**, IRE1/XBP1s target gene set used in **Fig. 2B,C**, and the PERK/ATF4 gene set used in **Fig. 2B**. Full RNAseq is available at GEO (GSE353599).

**Table S2. RNAseq analysis of differentiating T_H_17 cells treated for 24 h with IXA4 (10 µM) and/or STF-080310 (64 µM).** This excel file contains 5 individual spreadsheets including the normalized count matrix (cpm), Gene set Enrichment Analysis (GSEA) used in **Fig. 3A**, IRE1/XBP1s target gene set used in **Fig. S3B,C**, the PERK/ATF4 gene set used in **Fig. S3B,C**, and cholesterol homeostasis gene set used in **Fig. 3B,C**, Full RNAseq is available at GEO (GSE353599).

**Table S3. RNAseq analysis of differentiating T_H_17 cells treated for 24 h with IXA4 (10 µM) and/or MKC-3946 (10 µM).** This excel file contains 4 individual spreadsheets including the normalized count matrix (cpm), IRE1/XBP1s target gene set used in **Fig. S3D,E**, the PERK/ATF4 gene set used in **Fig. S3D,E**, and cholesterol homeostasis gene set used in **Fig. 3D**, Full RNAseq is available at GEO (GSE353599).

**Table S4. RNAseq analysis of differentiating T_H_17 cells treated for 6 h with IXA4 (10 µM) and/or desmosterol (10 µM)** This excel file contains 2 individual spreadsheets including the normalized count matrix (cpm) and the IRE1/XBP1s target gene set used in **Fig. S4E**. Full RNAseq is available at GEO (GSE353599).

**Table S5. RNAseq analysis of differentiating T_H_17 cells treated for 6 h with IXA4 (10 µM) or IXA62 (10 µM).** This excel file contains 3 individual spreadsheets including the normalized count matrix (cpm), IRE1/XBP1s target gene set used in **Fig. 5A,C**, and the PERK/ATF4 gene set used in **Fig. 5A,C**. Full RNAseq is available at GEO (GSE353599).

---

**Figure S1 (Supplement to Figure 1).**
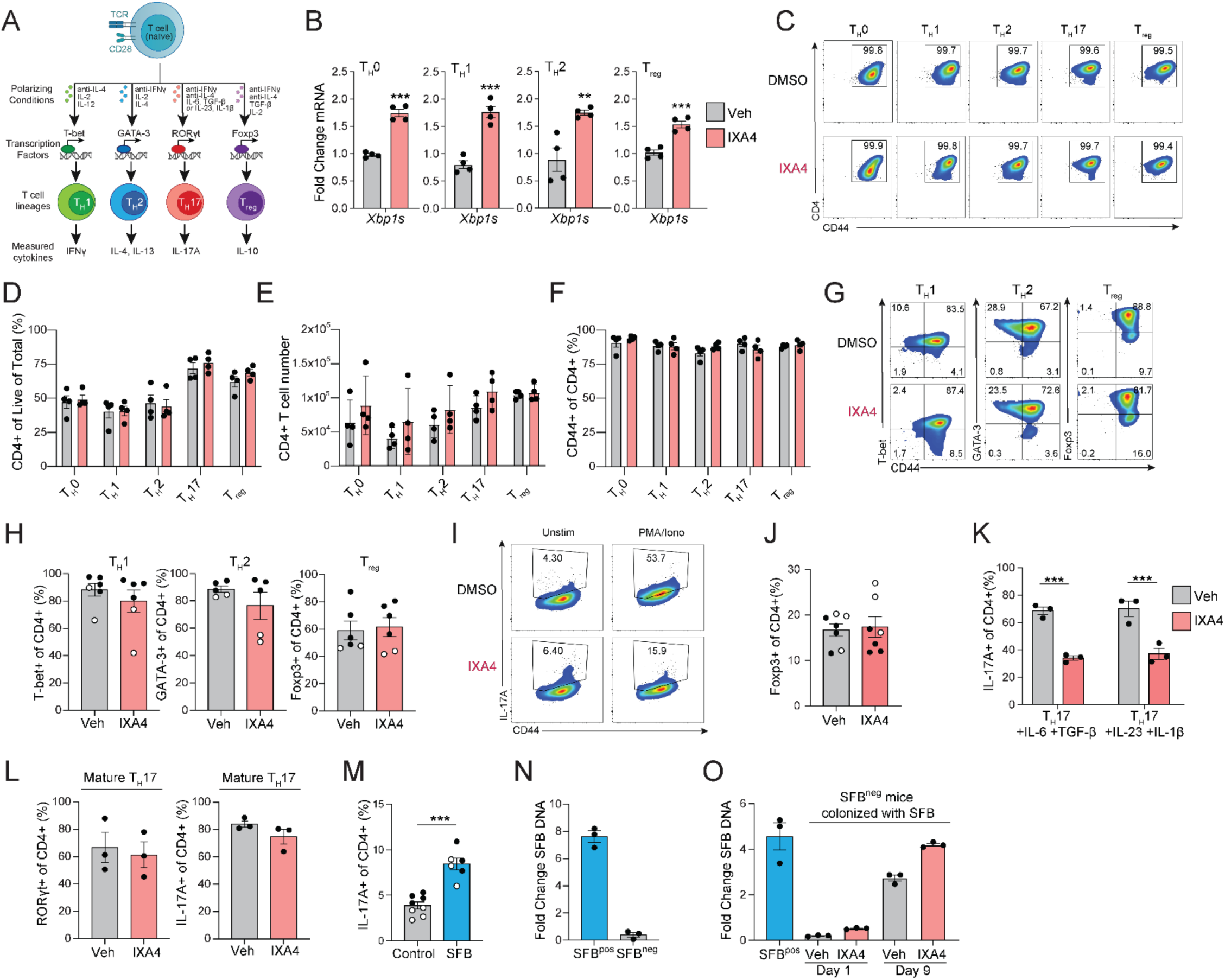
The IRE1/XBP1s activator IXA4 selectively impairs *ex vivo* and *in vivo* differentiation of CD4^+^ T_H_17 cells. **A.** Schematic of CD4^+^ T helper lineage specification. Naive CD4^+^ T cells activated through TCR and CD28 engagement differentiate into T_H_1, T_H_2, T_H_17, or T_reg_ lineages under the indicated polarizing conditions. Figure adapted from ref ^93^. **B.** Expression, measured by RT-qPCR, of *Xbp1s* transcript levels in CD4^+^ T cells polarized under T_H_1, T_H_2, T_H_17, or T_reg_ conditions for 24 h and then treated for 6 h with IXA4 (10 μM) or vehicle (DMSO). Error bars show SEM for n=4 biological replicates. **C.** Representative flow cytometry plots of CD4 and CD44 staining in CD4^+^ T cells polarized under the indicated conditions in the presence or absence of IXA4 (10 µM). **D,E.** Percent viable (**D**) and live cell count (**E**), measured by ZombieNIR staining, of CD4^+^ T cells polarized under the indicated conditions in the presence or absence of IXA4 (10 µM). Error bars show SEM for n=4 biological replicates. **F.** Percent CD4^+^ CD44^+^ cells in CD4^+^ T cells polarized under the indicated conditions in the presence or absence of IXA4 (10 µM). Error bars show SEM for n=4 biological replicates. **G,H.** Representative flow cytometry plots and quantification of T-bet (T_H_1), GATA-3 (T_H_2), or Foxp3 (T_reg_) staining versus CD44 staining in CD4^+^ T cells polarized under the indicated conditions in the presence or absence of IXA4 (10 μM). Error bars in (**H**) show SEM for n=6 biological replicates across two independent experiments. **I.** Representative flow cytometry plots of IL-17A staining in CD4^+^ T cells polarized in T_H_17 conditions in the absence or presence of IXA4 (10 µM). **J.** Percent Foxp3^+^ cells in CD4^+^ T cells polarized in T_H_17 conditions in the presence or absence of IXA4 (10 µM). Error bars show SEM for n=7 replicates across two independent experiments. **K.** Percent IL-17A^+^ CD4^+^ T cells polarized under standard T_H_17 conditions (IL-6, TGF-β) or alternative T_H_17 conditions (IL-23, IL-1β) in the presence or absence of IXA4 (10 µM). Error bars show SEM for n=3 replicates. **L.** Percent RORγt^+^ and IL-17A^+^ cells in CD4^+^ T cells polarized for 72 h in T_H_17 conditions and then treated for an additional 48 h with IXA4 (10 µM) or vehicle. Error bars show SEM for n=3 replicates. **M.** Percent IL-17A^+^ CD4^+^ T cells in Peyer’s patches of SFB^neg^ mice daily administered SFB^pos^ fecal slurry by oral gavage. Error bars show SEM for n=8 replicates across two independent experiments. **N**. SFB DNA, measured by qPCR and normalized to SFB^neg^, in feces from SFB^pos^ and SFB^neg^ mice. Error bars show SEM for n=3 biological replicates. **O**. SFB DNA, measured by qPCR and normalized to SFB^neg^, in feces from mice administered fecal slurry and/or IXA4 (50 mg/kg) daily by oral gavage, assessed at day 1 and day 9 of treatment. Feces from SFB^pos^ mice are shown as a reference. Error bars show SEM for n=3 biological replicates. **p<0.01, ***p<0.001 for two-tailed unpaired t-test (**B, H, M**) or 2-way ANOVA (**K**).

**Figure S2 (Supplement to Figure 2).**
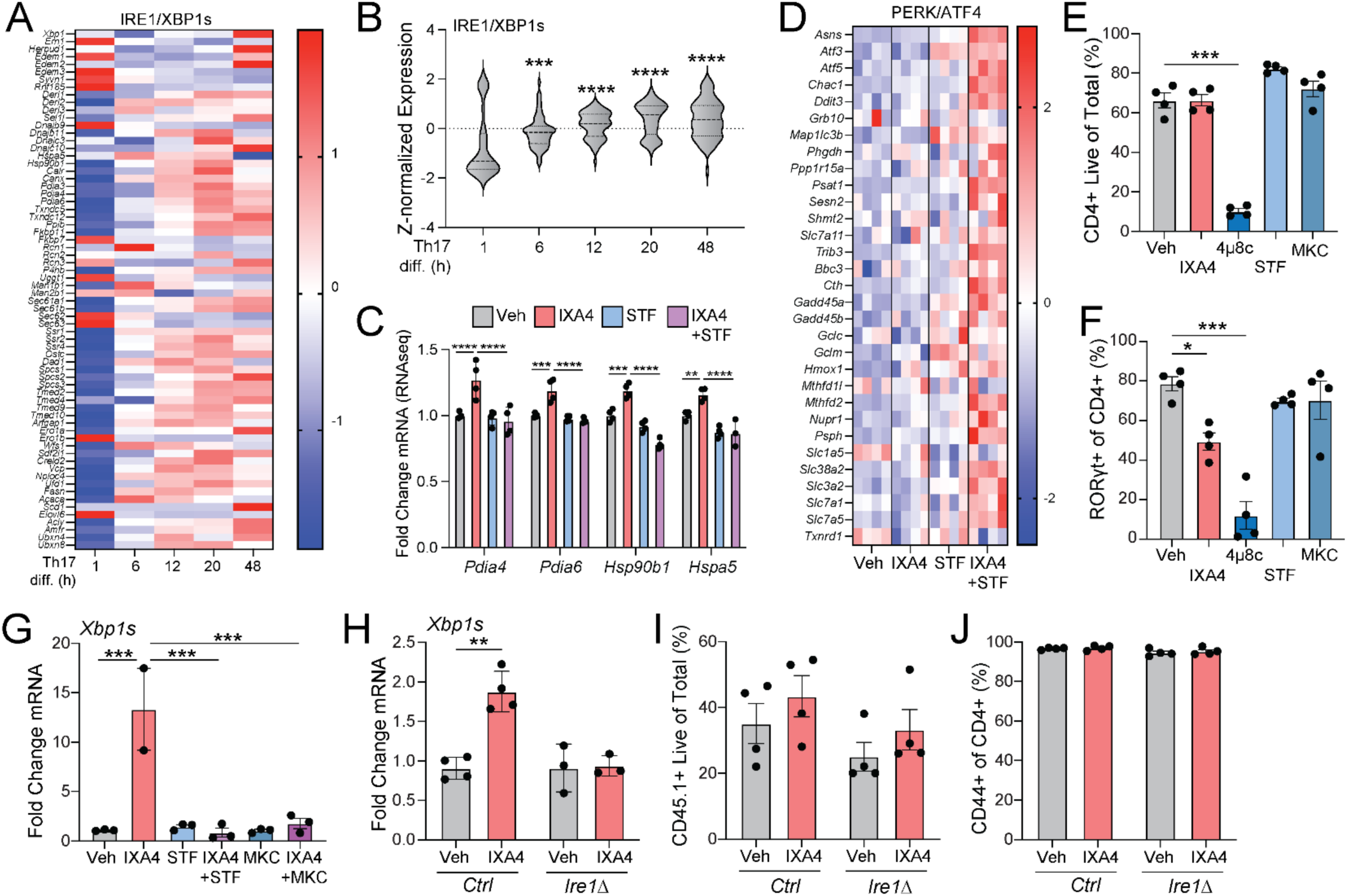
IXA4-mediated reductions in T_H_17 differentiation require IRE1 activity. **A.** Heatmap of IRE1/XBP1s target gene expression across a time course of T_H_17 differentiation (1, 6, 12, 20, and 48 h). Data from ref^61^. **B.** Z-normalized expression of the IRE1/XBP1s target gene set described in (**A**) across the T_H_17 differentiation time course. **C.** Expression, measured by RNAseq, of the IRE1/XBP1s target genes *Pdia4, Pdia6, Hsp90b1,* and *Hspa5* in differentiating T_H_17 cells treated for 6 h with IXA4 (10 µM) and/or STF-083010 (64 µM). Error bars show SEM for n=4 biological replicates. **D.** Heat map of Z-normalized gene expression for PERK/ATF4 target genes from RNAseq data described in Fig. 2B. **E.** Viability, measured by ZombieNIR, of CD4^+^ T cells polarized under T_H_17 conditions treated with IXA4 (10 µM), 4µ8c (32 µM), STF-083010 (64 µM), or MKC-3946 (10 µM). Error bars show SEM for n=4 biological replicates. **F.** Quantification of RORγt^+^ cells in differentiating T_H_17 cells treated as described in (**E**). Error bars show SEM for n=4 biological replicates. **G.** *Xbp1s* mRNA, measured by RT-qPCR, in differentiating T_H_17 cells treated with IXA4 (10 µM), STF-083010 (64 µM), and/or MKC-3946 (10 µM) for 24 h. Error bars show SEM for n=2-3 biological replicates. **H.** *Xbp1s* mRNA, measured by RT-qPCR, in CD4^+^ T cells isolated from *Ctrl* or *Ire1Δ* mice, polarized under T_H_17 conditions, and treated with IXA4 (10 µM) or vehicle for 24 h. Error bars show SEM for n=4 biological replicates **I.** Viability, measured by ZombieNIR, of *Ctrl* or *Ire1Δ* CD4^+^ T cells polarized under T_H_17 conditions and treated with IXA4 (10 µM) or vehicle. Error bars show SEM for n=4 biological replicates. **J.** Activation status, measured by CD4 and CD44 staining, of cells as described in (**I**). Error bars show SEM for n=4 biological replicates. *p < 0.05, **p < 0.01, ***p < 0.001, ****p < 0.0001 for one-way ANOVA (**B, E, F, G**), two-way ANOVA (**C**), or unpaired t-test (**H**).

**Figure S3 (Supplement to Figure 3).**
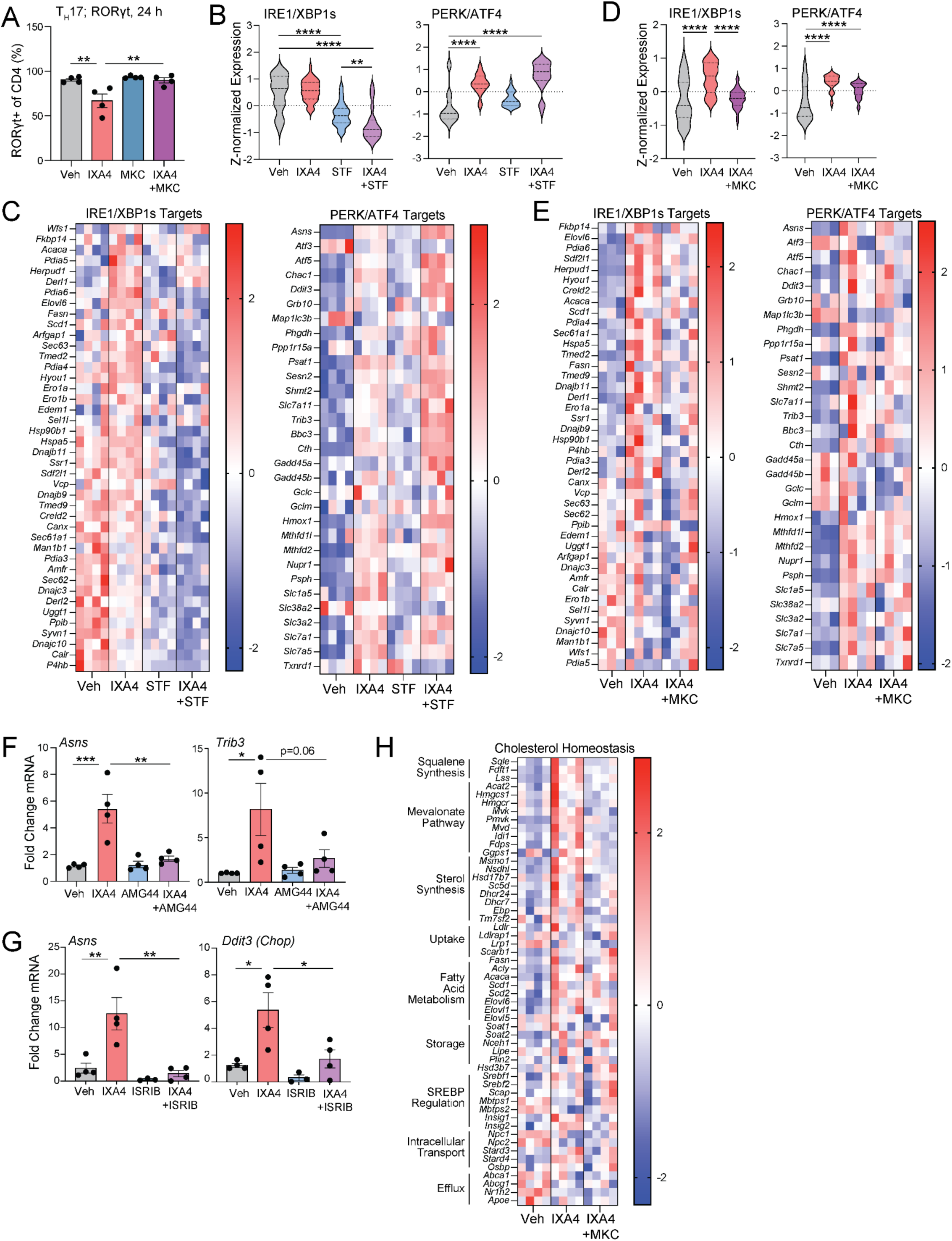
IXA4-mediated IRE1 activation depletes intracellular cholesterol in differentiating T_H_17 cells. **A.** Percent RORγt^+^ cells in CD4^+^ T cells polarized under T_H_17 conditions for 24 h and then treated an additional 24 h with IXA4 (10 µM) +/-MKC-3946 (10 µM). **B.** Z-normalized expression of gene sets comprising IRE1/XBP1s target genes (left) and PERK/ATF4 target genes (right) in differentiating T_H_17 cells treated for 24 h with IXA4 (10 µM) +/-STF-083010 (64 µM). Full gene sets are shown in **Table S2**. **C.** Heatmaps of IRE1/XBP1s target genes (left) and PERK/ATF4 target gene (right) expression across the conditions described in (**B**). Each column represents an individual biological replicate. **D.** Z-normalized expression of IRE1/XBP1s and PERK/ATF4 target gene sets in differentiating T_H_17 cells treated for 24 h with IXA4 (10 µM) +/-MKC-3946 (10 µM). Full gene sets are shown in **Table S3**. **E.** Heatmaps of XBP1s target genes (left) and PERK/ATF4 target genes (right) expression across the conditions described in panel (**D**). **F**. *Asns* and *Trib3* mRNA levels, measured by RT-qPCR, in differentiating T_H_17 cells treated for 48 h with IXA4 (10 µM) in the absence or presence of the PERK kinase inhibitor AMG44 (10 µM). **G.** *Asns* and *Ddit3* mRNA levels, measured by RT-qPCR, in differentiating T_H_17 cells treated for 48 h with IXA4 (10 µM) in the absence or presence of the PERK signaling inhibitor ISRIB (250 nM). **H.** Heatmap of cholesterol homeostasis gene expression in differentiating T_H_17 cells treated for 24 h with IXA4 (10 µM) +/-MKC-3946 (10 µM). Each column represents a biological replicate. *p < 0.05, **p < 0.01, ****p < 0.0001 for one-way ANOVA.

**Figure S4 (Supplement to Figure 4).**
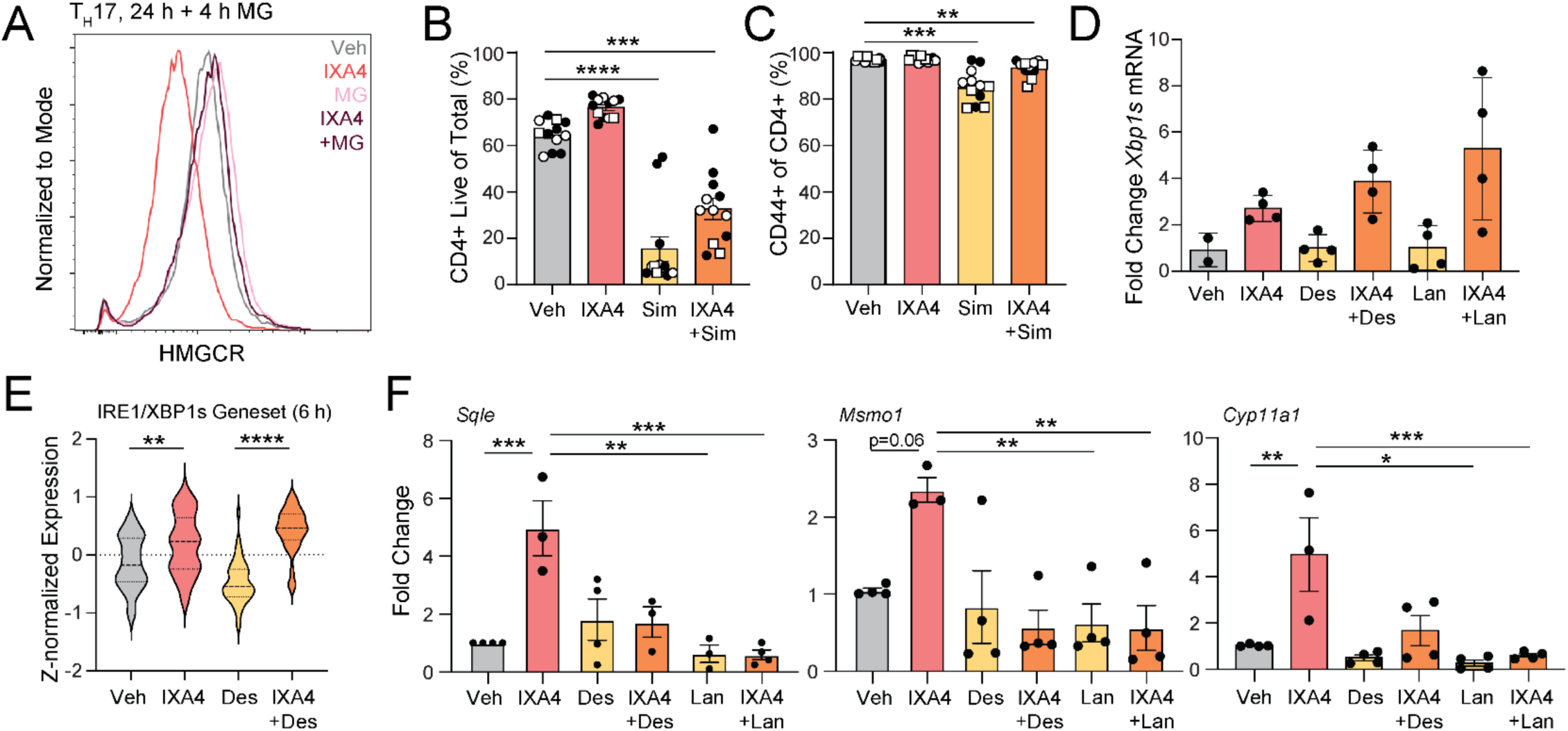
Co-treatment with cholesterol precursors prevents IXA4-mediated reductions in T_H_17 differentiation. **A.** Representative histograms, measured by flow cytometry, for HMGCR staining in CD4^+^ T cells polarized under T_H_17 conditions and treated for 24 h with IXA4 (10 µM) and then an additional 4 h with MG132. B, C. Viability, measured by ZombieNIR (B) and activation status, measured by CD4 and CD44 staining (C), of CD4^+^ T cells polarized under T_H_17 conditions and treated with IXA4 (10 µM) and/or simvastatin (1 µM). Error bars show SEM for n=12 replicates across 3 independent experiments. D. *Xbp1s* mRNA levels, measured by RT-qPCR, in differentiating T_H_17 cells treated for 6 h with IXA4 (10 µM), desmosterol (Des, 10 µM), and/or lanosterol (Lan, 10 µM). Error bars show SEM for n=3-4 replicates. E. Z-normalized expression of the IRE1/XBP1s target gene set in differentiating T_H_17 cells treated for 6 h with IXA4 (10 µM) in the absence or presence of desmosterol (10 µM). Full gene sets are described in Table S4. F. Expression of the cholesterol biosynthesis genes *Sqle, Msmo1*, and *Cyp11a1* measured by RT-qPCR in differentiating T_H_17 cells treated for 48 h with IXA4 (10 µM) in the absence or presence of desmosterol (10 µM) or lanosterol (10 µM). Error bars show SEM for n=3-4 replicates. *p < 0.05, **p < 0.01, ***p < 0.001, ****p < 0.0001 for one-way ANOVA.

**Figure S5 (Supplement to Figure 5).**
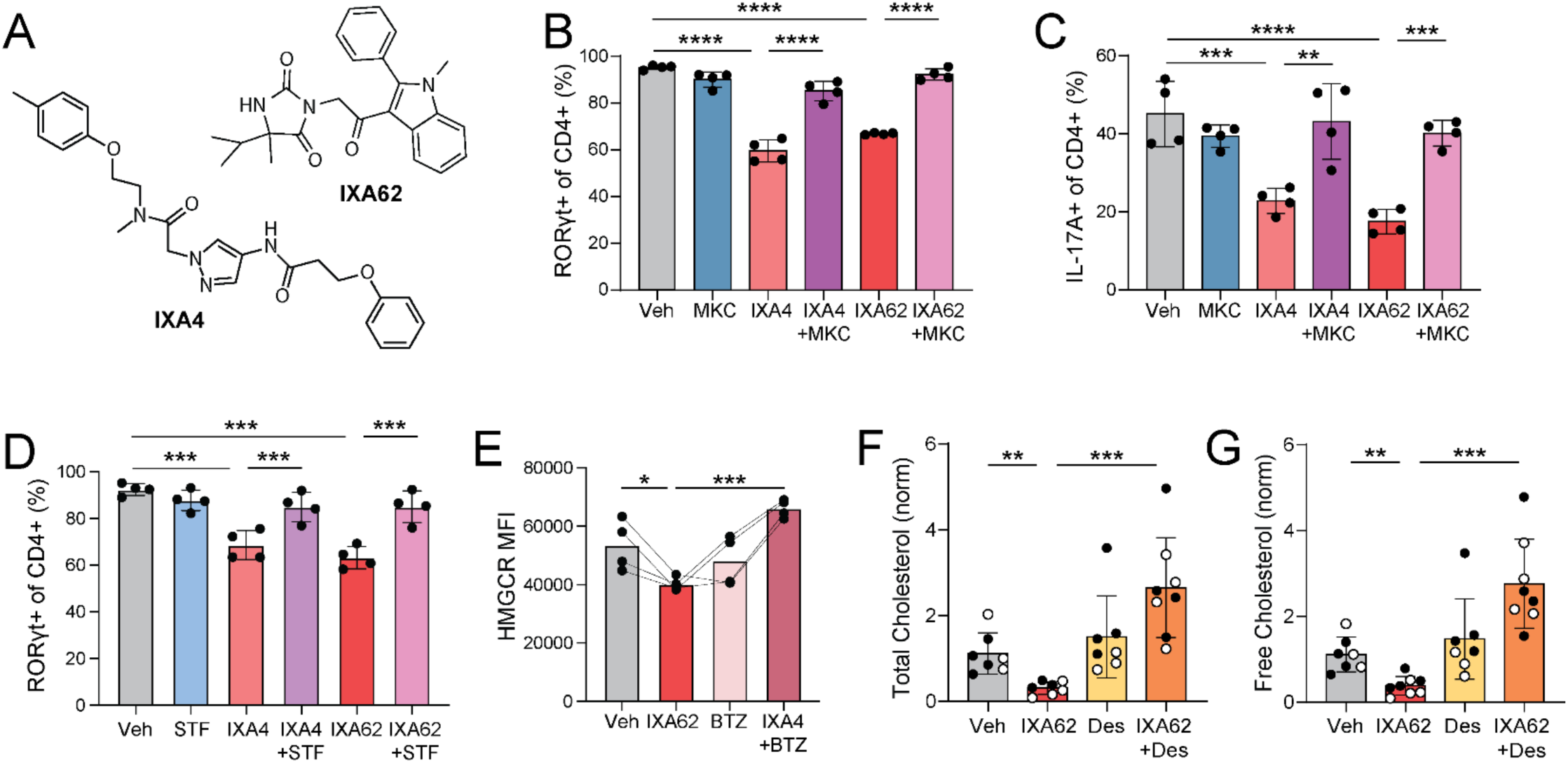
IXA62-mediated IRE1 activation reduces T_H_17 differentiation by rewiring cholesterol metabolism. **A.** Chemical structures of IXA4 and IXA62. B, C. Percent RORγt⁺ (B) and IL-17A⁺ (C) cells in CD4⁺ T cells polarized under T_H_17 conditions and treated with IXA4 (10 µM) or IXA62 (10 µM) in the absence or presence of MKC-3946 (10 µM). Error bars show SEM for n=4 replicates. D. Percent RORγt⁺ cells in differentiating T_H_17 cells treated with IXA4 (10 µM), IXA62 (10 µM), and/or STF-083010 (64 µM). Error bars show SEM for n=4 replicates. E. HMGCR levels measured by flow cytometry in differentiating T_H_17 cells treated with IXA62 (10 µM) for 24 h and then an additional 4 h with bortezomib (1 nM). Error bars show SEM for n=4 replicates. F, G. Normalized total (F) and free (G) cholesterol in differentiating T_H_17 cells treated with IXA62 (10 µM) in the absence or presence of desmosterol (10 µM). Error bars show SEM for n=7-8 replicates across 2 independent experiments. \**p* < 0.05, \*\**p* < 0.01, \*\*\**p* < 0.001, \*\*\*\**p* < 0.0001 for one-way ANOVA.

## EXTENDED DATA FIGURES

**Extended Data Figure 1.**
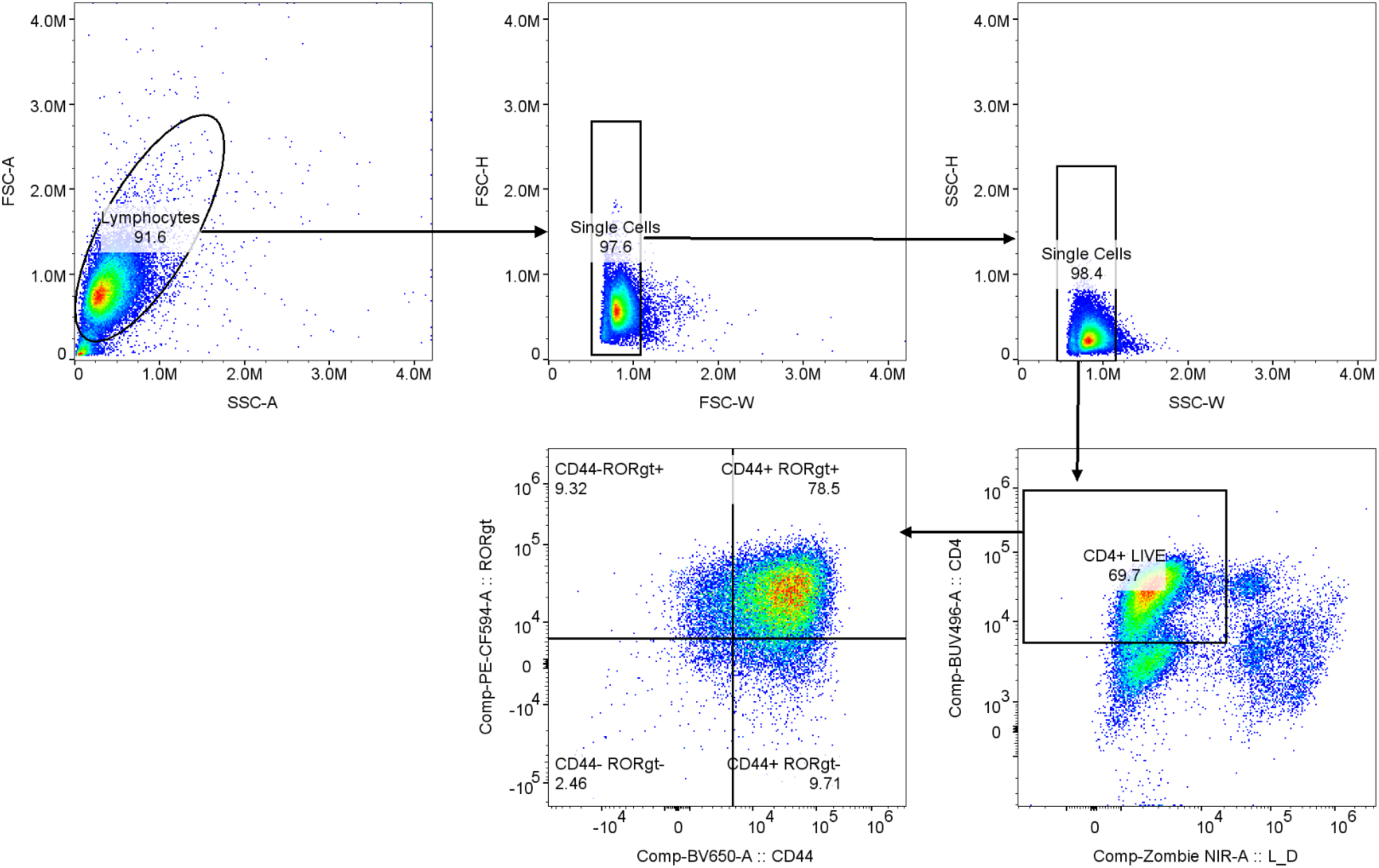
Gating Strategy for RORγt in CD4^+^ T_H_17 cells (Relevant to Figure 1).

**Extended Data Figure 2.**
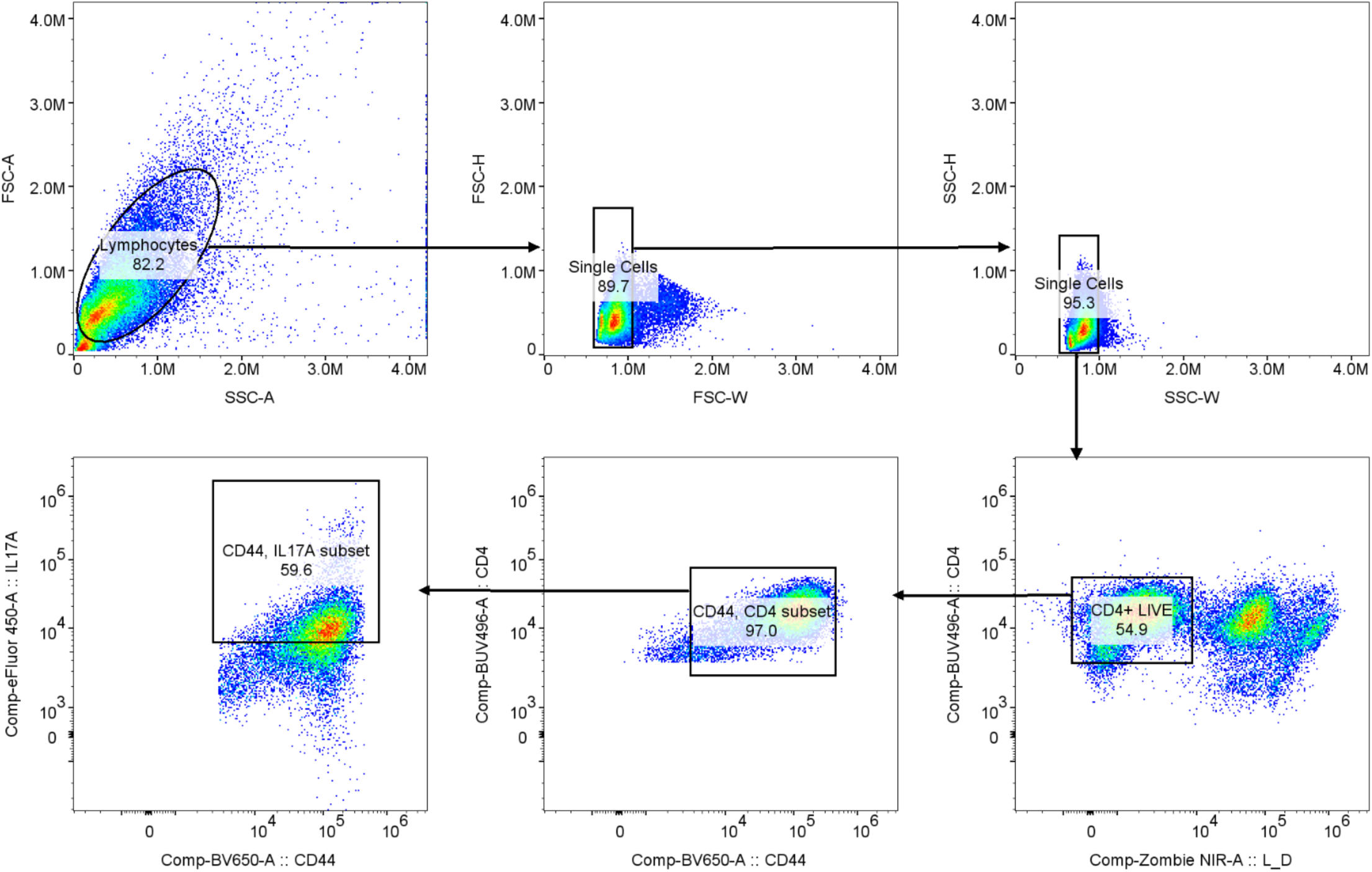
Gating Strategy for IL-17A in CD4^+^ T_H_17 cells (Relevant to Figure 1).

**Extended Data Figure 3.**
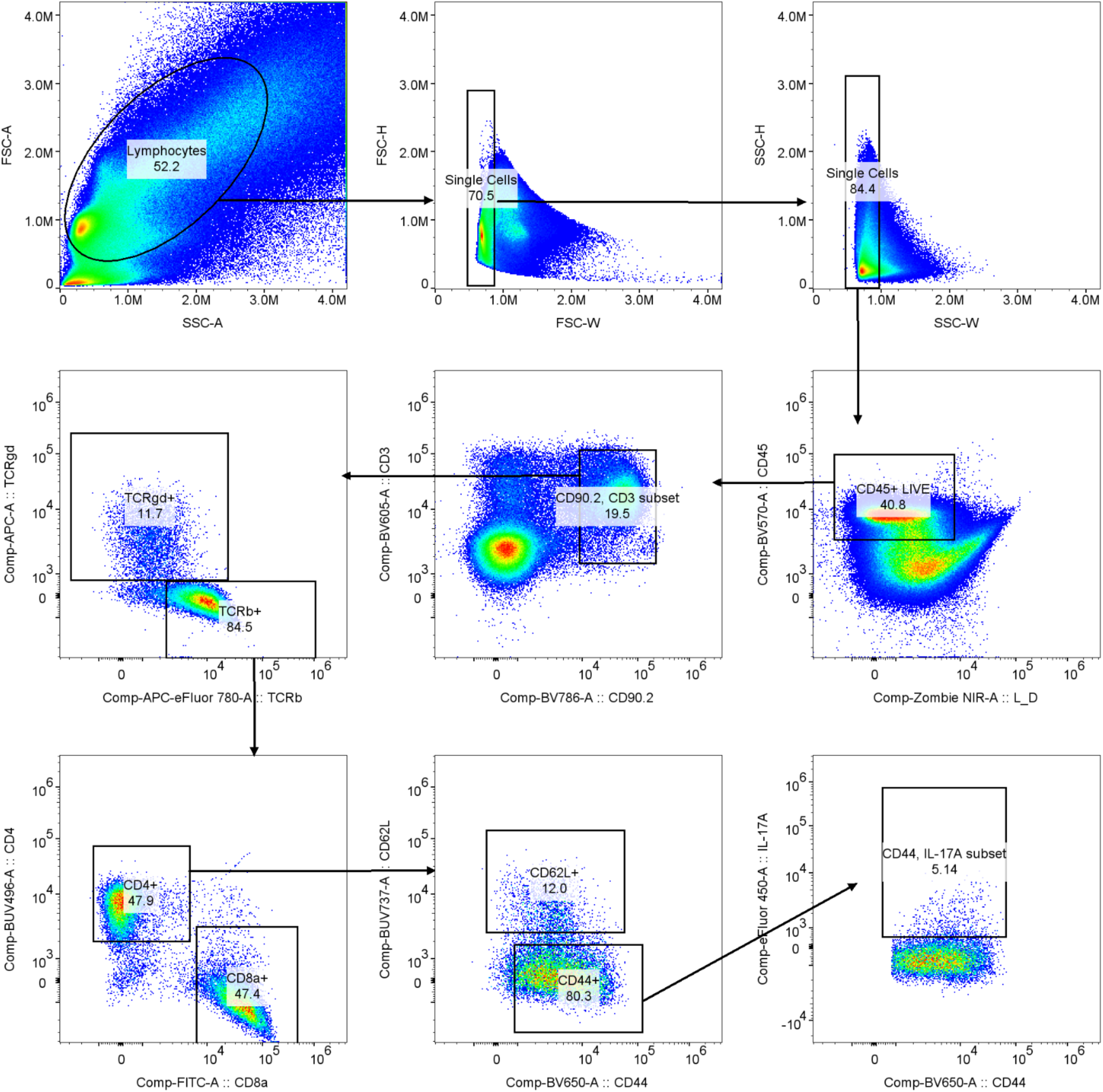
Gating Strategy for CD4^+^ IL-17A^+^ T cells in Peyer’s Patches. (Relevant to Figure 1).

**Extended Data Figure 4.**
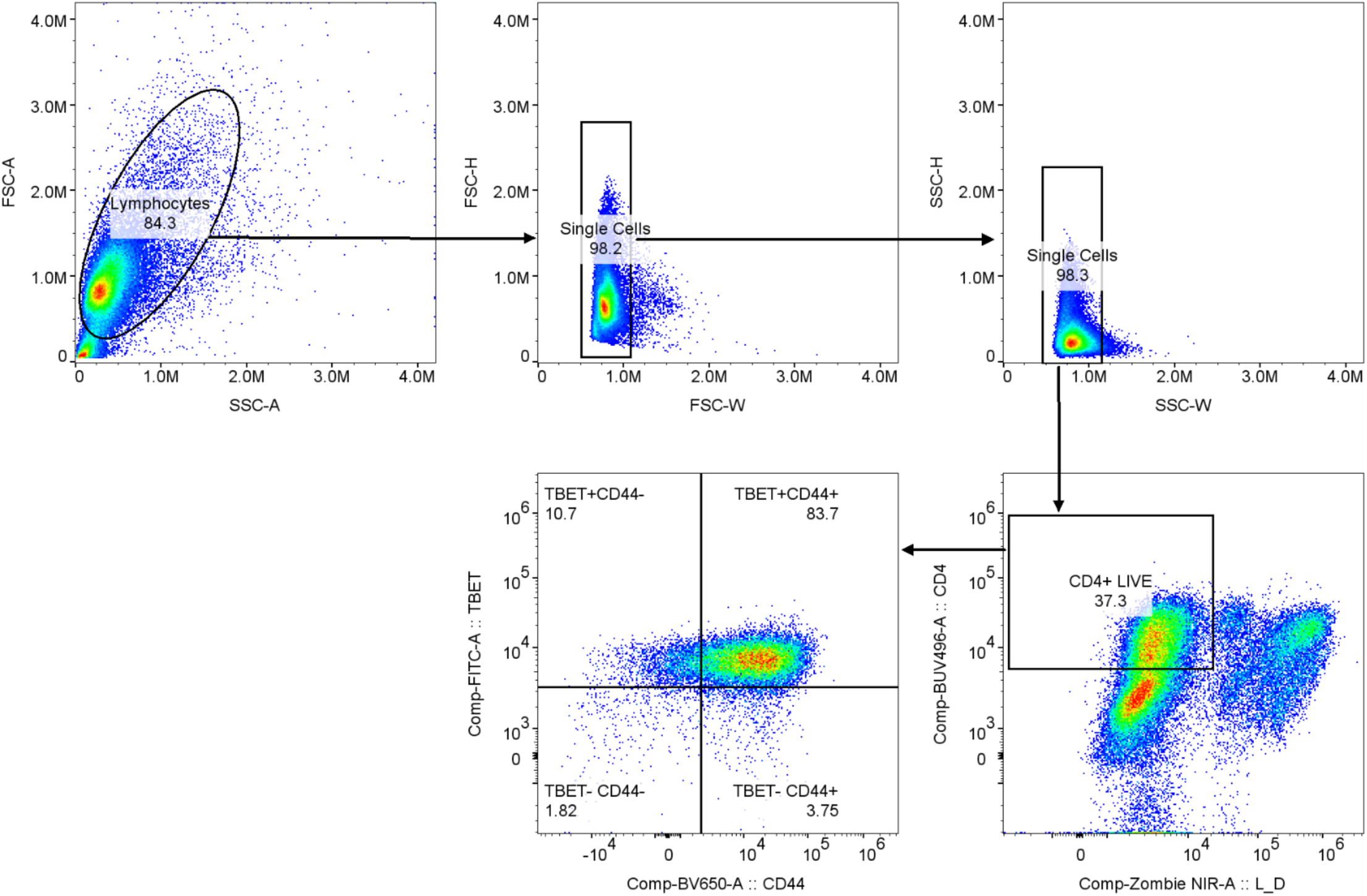
Gating Strategy for T-bet in CD4^+^ T_H_1 cells (Relevant to Figure S1).

**Extended Data Figure 5.**
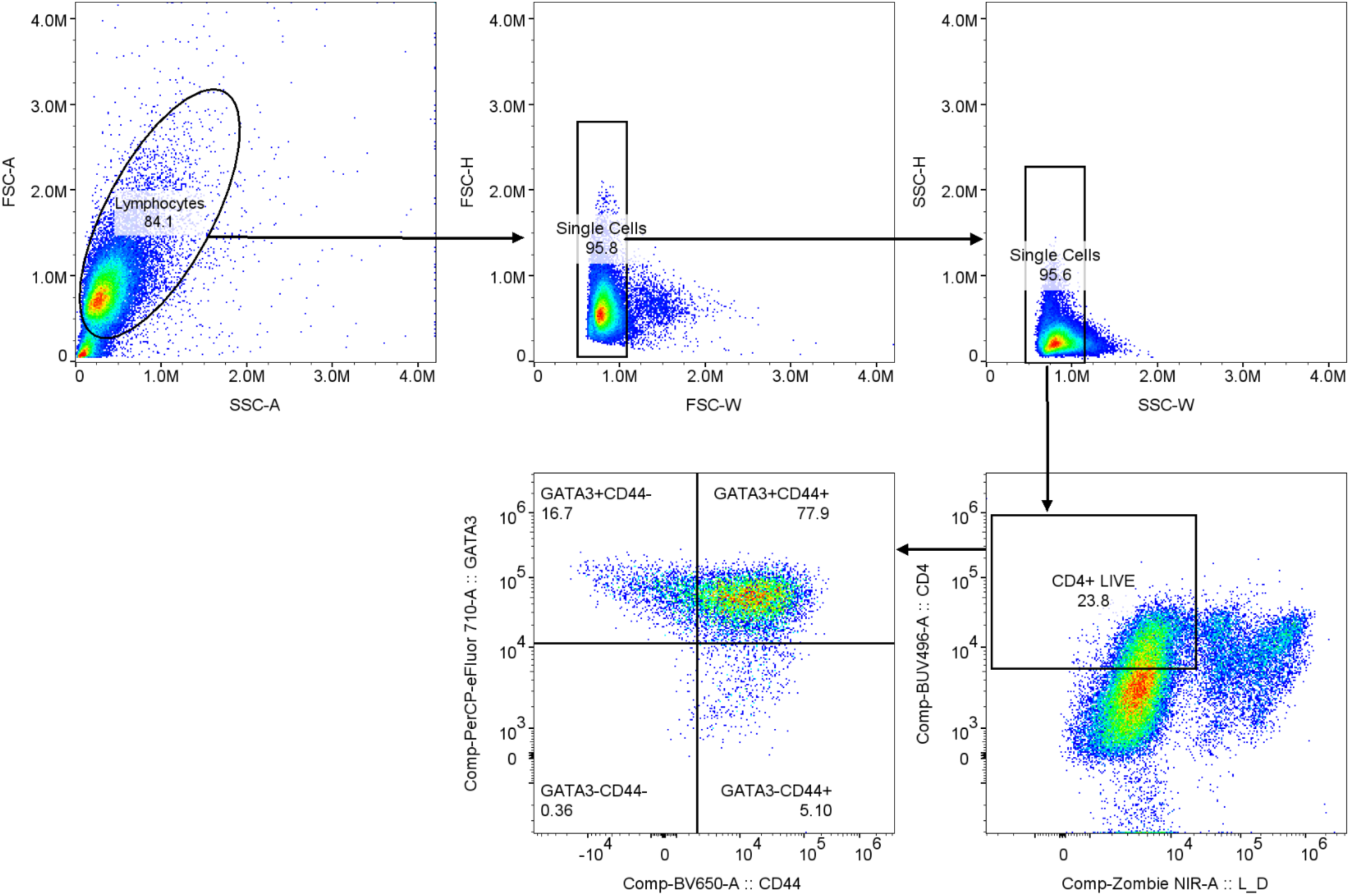
Gating Strategy for GATA-3 in CD4^+^ T_H_2 cells (Relevant to Figure S1).

**Extended Data Figure 6.**
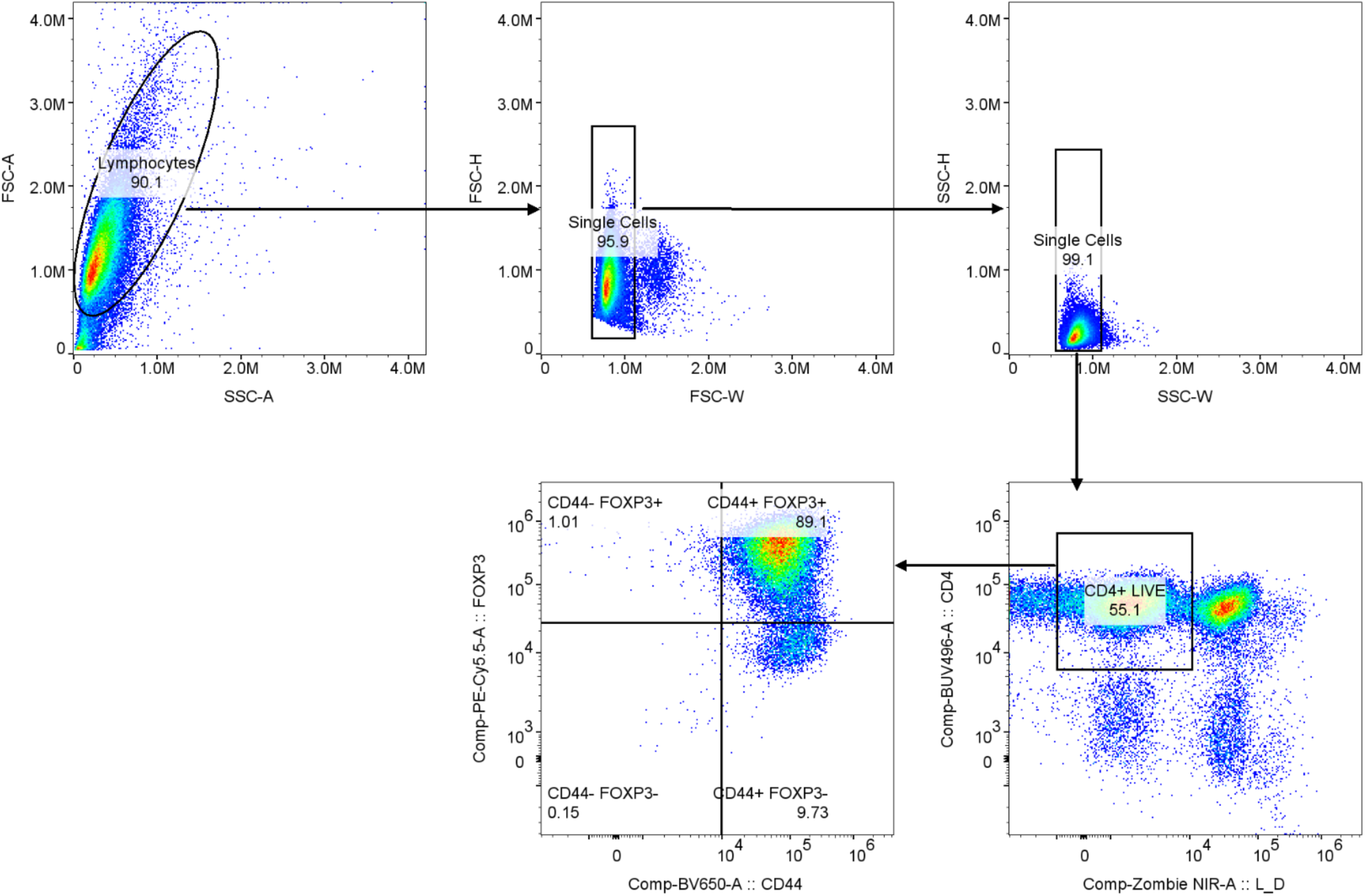
Gating Strategy for Foxp3 in CD4^+^ T_reg_ cells (Relevant to Figure S1).

